# Discovery of small molecule probes targeting the 5′ stem-loop in the yeast U4/U6 snRNA

**DOI:** 10.1101/2025.10.10.681657

**Authors:** Mo Yang, Shaifaly Parmar, Sumirtha Balaratnam, Desta D. Bume, Peri R. Prestwood, Wojciech K. Kasprzak, John S. Schneekloth

## Abstract

The spliceosome is a large ribonucleoprotein complex that regulates pre-mRNA splicing and has been an intriguing target for drug discovery. Essential to the assembly of the spliceosome are the small nuclear RNAs (snRNAs), which form RNA-RNA and RNA-protein interactions in the intact spliceosome and during its assembly. Here, we study the yeast U4/U6 snRNA assembly and report the rapid discovery of small molecule K-turn ligands via parallel small molecule microarray (SMM) screening of multiple related RNA constructs. For hit validation, biophysical analyses were conducted to confirm the binding and effects on thermodynamic stability and of RNA structure. One analog of the hit molecule (**22**) exhibited improved affinity towards the yeast U4/U6 snRNA (K_D_ = 3.9 ± 2.2 µM). The specific interaction between 22 and the K-turn region was studied experimentally using deltaSHAPE and *in silico* with MD simulations. Moreover, this molecule was found to inhibit the binding of the U4 to Snu13 in biochemical assays (IC_50_ = 3.2 ± 0.4 µM). This work reports new ligands for the U4 snRNA and reveals that a multiplexed, structure-based approach can be used to identify small molecules that bind to specific regions of complex RNAs.

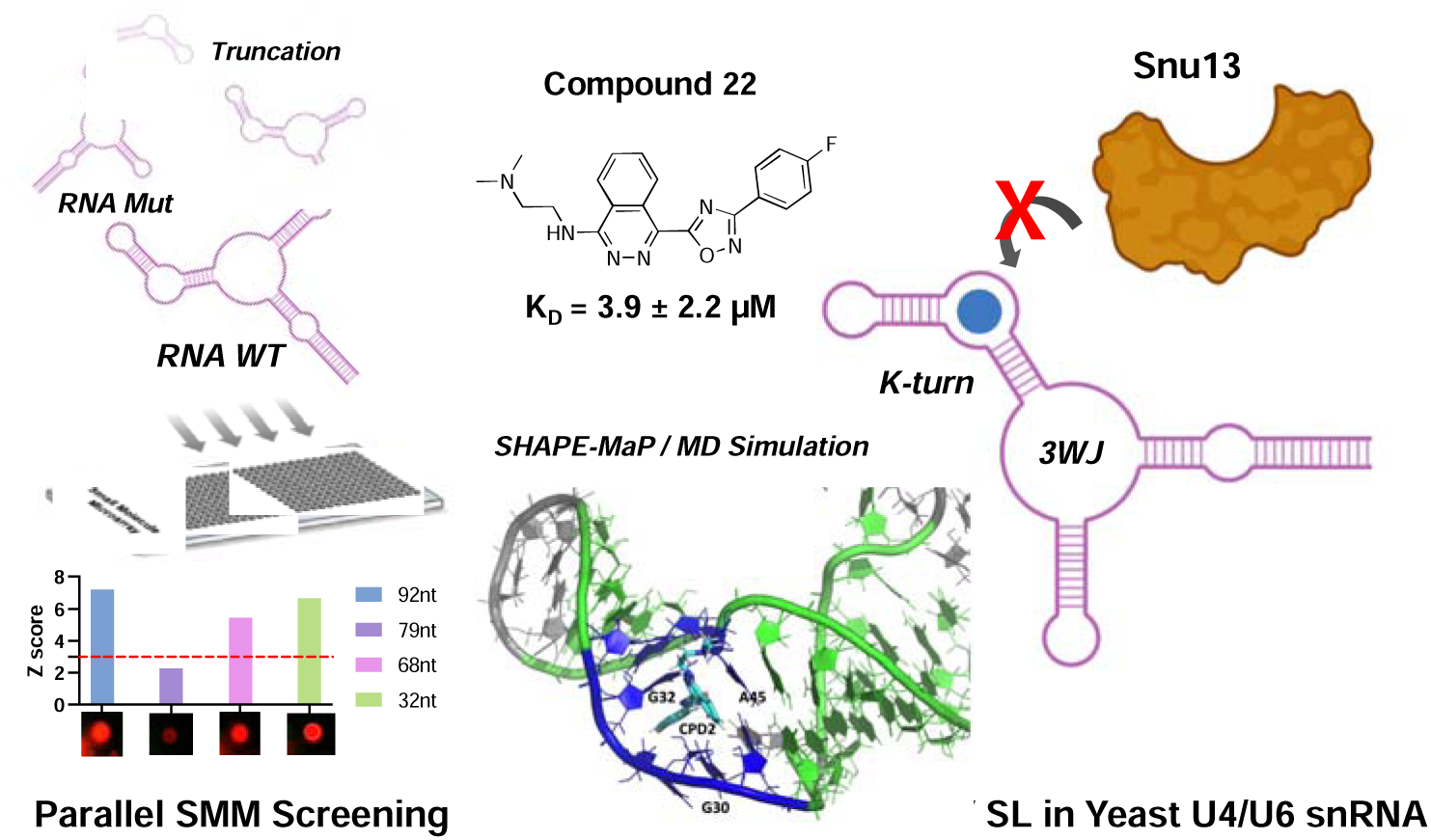

## INTRODUCTION

The spliceosome is a large ribonucleoprotein complex that governs pre-mRNA splicing.^1, 2^ Splicing is a key component of gene expression and contributes broadly to transcriptomic and proteomic diversity as well as many diseases including cancer.^3, 4^ The disease relevance, high structural complexity of the spliceosome (comparable to the ribosome), and dynamic nature of its structure (assembled and disassembled during each splicing event) all contribute to interest in studying small molecules that impact spliceosome assembly and function. To date most of the developed molecules targeting the spliceosome focus on protein targets, such as SF3B inhibitors (e.g., Spliceostatin A, H3B-8800) and CDK7 inhibitors (e.g., SY-5609, CT7001) for cancer.^5, 6^ However, molecules targeting the small nuclear RNAs (snRNAs) that make up the RNA component of the spliceosome are also intriguing.

RNA as a target for small molecules has been attracting increased attention from both academia and the pharmaceutical industry due to its great potential for development of novel therapeutics in challenging areas including cancer, spinal muscular atrophy (SMA), Huntington’s disease and cystic fibrosis, etc.^4, 7–12^ Recently, Risdiplam, a small molecule that targets survival motor neuron-2 (SMN2) splicing by binding to the U1 snRNP, has been approved by the FDA for the treatment of SMA.^13, 14^ Other splicing modulators are also in development such as PTC518 (Phase II clinical trials) for Huntington’s disease,^15^ and compounds targeting MAPT exon 10 for Frontotemporal dementia with parkinsonism (FTDP).^16^ This encouraging progress highlights splicing as a key target in multiple diseases.^17^

Targeting key components of the spliceosome itself provide great opportunities to develop novel chemical probes and potential therapeutics.^18–20^ The assembly of the spliceosome is highly complex, dynamic, and dependent on multiple proteins and snRNAs. During spliceosome assembly, the U4 and U6 snRNAs interact through base pairing and recruit multiple proteins, during the assembly of the pre-catalytic spliceosome.^1, 21^ The U4/U6 di-snRNA assembly also plays an essential role in recruiting downstream proteins to form U4/U6.U5 tri-snRNPs.^22^ The U4 snRNA contains a kink turn (K-turn) motif embedded in the 5′ stem-loop (5′ SL) that binds to the small nuclear ribonucleoprotein 13 (Snu13) in yeast.^23^ Recent studies on the ligandability of RNA structures have reported that RNAs that fold into complex structures can contain hydrophobic pockets suitable for small molecule binding.^24, 25^ We demonstrate that the U4 K-turn contains a hydrophobic pocket that could potentially be targeted by small-molecule probes. Several structural studies have revealed that protein binding alters the conformation of U4 K-turn, suggesting that ligand binding might impact this process.^26–28^ For example, previous efforts have shown that topotecan, a topoisomerase I inhibitor and analog of the natural product camptothecin that binds broadly to nucleic acids, disrupts the binding between NHP2L1 (Snu13) and U4 5′ SL by binding to RNA.^29^

To date, significant efforts have been made to target RNA with small molecules.^30^ However, it remains challenging to identify and develop small molecule RNA ligands,^8, 31^ and new approaches for ligand discovery are still needed. Small molecule microarray (SMM) technology has been developed as a powerful tool for target-based high-throughput screening in a cost-effective manner.^32^ Recently, SMMs focusing on RNA targets have been utilized to realize drug-like compound discovery as well as RNA-ligand recognition studies.^33^ A key advantage of the SMM approach is the ability to rapidly counter screen against multiple other RNA constructs, providing information about selectivity early in the discovery process.

Herein, we report small molecule probes that target the specific pocket region of yeast U4/U6 snRNA. A targeted SMM screening strategy was developed by using wild type (WT), mutated (mut), and truncated U4/U6 RNA constructs, facilitating the rapid discovery and prioritization of molecules that specifically bind the K-turn region of the U4. Further biophysical studies confirmed that the molecules directly bind and stabilize the intended RNA structure, in agreement with the molecular dynamics (MD) simulation results. An improved analog was found to block the binding between the 5′ SL RNA and Snu13 protein. To our knowledge, this is the first example of compounds that bind the yeast U4/U6 snRNA at the precatalytic stage of splicing. This study provides new perspectives on targeting the spliceosome as well as modulating protein-RNA interactions during RNA splicing.

## METHODS

### General Information

U4/U6 snRNA constructs for high-throughput screening and biophysical experiments were chemically synthesized by Dharmacon (GE Healthcare). The sequences are summarized in **Supplementary Table S1**. The RNA constructs were purchased from Dharmacon with no labeling, or with biotin/Cy5 labeling at 5′ end. All buffers were prepared with RNase-free water (DEPC-treated) to avoid RNA degradation. Before use, the RNA was folded in folding buffer (11.25 mM KH_2_PO_4_/K_2_HPO_4_, pH 7.0, 310 mM KCl, 5 mM EDTA) at a concentration lower than 20 µM and annealed by heating at 90 for 5 min followed by snap cooling on ice. Then RNA was concentrated in spin columns (Merck Millipore, 3,000 Da cutoff) or diluted in other working buffers before future experiments. For hit validation and other biophysical/biochemical assays, the compounds and analogs (**1**-**22**) were purchased from ChemDiv, and **23**-**25** were synthesized in the lab (see Supplementary Information). All compounds were prepared as 10 mM stocks in DMSO.

### High-throughput Screening Using small molecule microarrays (SMMs)

For high-throughput screening using SMM, the microarray slides were prepared based on previously reported methods using a robotic microarray printer (Arrayjet, UK).^34^ Each slide contained ∼7,000 compounds with duplicate spots of each sample. To perform screening, the SMM slide surface was first blocked with 500 nM yeast total tRNA (Invitrogen) in screening buffer (1xPBS, pH 7.4, 0.005% Tween 20) for 2 h prior to target RNA incubation. Then, the 5′-Cy5-labeled U4/U6 RNA construct was folded as above described, followed by buffer exchange to screening buffer. After annealing, the RNA was diluted to 50 nM in screening buffer and incubated with the microarray for another 2 h. After that, the glass slide was gently washed with PBST, PBS, and ddH_2_O three times in each buffer solution and dried in a centrifuge at 1,700 g for 2 min. Finally, the microarray was imaged with an InnoScan 1100 AL fluorescence scanner (Innopsys, France) at 635 nm and the hits were identified based on the quantified fluorescent intensity of each spot.

### Surface Plasmon Resonance (SPR)

SPR experiments were carried out using an SPR optical biosensor (BIAcore 3000, Cytiva, USA) based on reported methods with minor modifications.^35^ To avoid RNA degradation during long-time injections, the whole microfluidic system was washed by priming 50% RNase Zap buffer and RNase-free water one day before the experiments. Then, by using running buffer (1xPBS, pH7.4, 0.005% Tween 20, 5% DMSO) at a flow rate of 5 µL/min, the binding assays were performed. In brief, a CM5 chip surface coated with carboxylated dextran was activated by standard EDC/NHS amine-coupling protocol and then conjugated with streptavidin (200 ng/mL in 10 mM sodium acetate buffer, pH 4.5) for 30 min, resulting in 4000∼6000 response units (RUs) signal. After streptavidin conjugation, the surface was deactivated (1 M ethanolamine, pH 8.5) for 10 min and regenerated (10 mM NaOH) for 2 min to remove unbound SA. Then, the folded U4/U6 RNA with biotin-label at 5′ end was prepared (according to the above-mentioned method) at 5 µM and injected in flow-cell (Fc) 2 for 30 min while Fc1 was kept blank as a reference. Then, the system continued flowing the running buffer until the baseline was stable before future steps.

To detect binding between small molecules and U4/U6 snRNA, a 50 µL compound solution was applied to both Fc1 and Fc2 channels simultaneously at a higher flow rate (25 µL/min). The binding curve contains 120 s association and 200 s dissociation. A pulse of 10 mM NaOH injection was used for regeneration if needed. To measure the dissociation constant (K_D_), titrations of compound solutions with a serial dilution were applied. The K_D_ value was then calculated by fitting the binding curves based on a steady-state model in BIAevaluation 4.0 software and plotted in GraphPad Prism 9.

### Microscale Thermophoresis (MST)

MST experiments were conducted using a Monolith NT.115 system (NanoTemper, Germany) as described previously.^36^ 5′-Cy5-labeled U4/U6 RNA was folded as above-mentioned and diluted to 100 nM (2x) in a working buffer. For RNA-small molecule interactions, compounds were prepared in the running buffer (with 10% DMSO) and mixed with RNA in a 1:1 (v/v) ratio, resulting in 50 nM RNA and 5% DMSO in the final solution. Then, MST scanning was performed in triplicate using a red LED channel to obtain the binding signals. Binding affinity (K_D_) was determined by fitting the concentration-dependent curve in the T-jump region according to the 1:1 Langmuir binding model (MO.Affinity Analysis v2.3).

For the protein-RNA inhibition assay, Snu13 protein was prepared in the running buffer (10 mM HEPES, pH 7.4, 150 mM NaCl, and 0.005% Tween 20) at different concentrations ranging from 0 µM to 5 µM. Meanwhile, a compound at a designated concentration was added to the annealed RNA solution. Then, the test solutions were prepared by 1:1 (v/v) mixing the protein and RNA-small molecule samples, followed by the standard MST procedure mentioned above.

### Circular Dichroism (CD)

All CD spectra were collected by a CD spectrometer (JASCO, USA). Unlabeled U4/U6 snRNA was folded (based on the above method, see SMM screening section) and prepared in a quartz cuvette at 5 µM, followed by triplicate scanning from 320 nm to 200 nm at an interval of 1 nm. To test the effect of the compound on RNA stability, a thermal melting assay was performed. In PBS buffer, folded U4/U6 snRNA was mixed with/without the compound at a designated concentration (5 µM RNA and 5% DMSO in the final solution). 300 µL of the RNA solution in a cuvette was gradually heated from 20 to 90 with an interval of 1 . To determine the melting temperature (T_m_), the peak at ∼260 nm was monitored in real time throughout the whole melting experiment. Then the temperature-dependent peak height was plotted. Finally, T_m_ was calculated based on the temperature corresponding to the peak of the first derivative curve using GraphPad Prism 9 software.

### Selective 2′-hydroxyl acylation analyzed by primer extension and mutational profiling (SHAPE-MaP) Based Structure Probing of U4U6 snRNA

SHAPE-MaP experiments were carried out based on previously reported protocols, and 2-aminopyridine-3-carboxylic acid imidazolide (2A3) was utilized as a SHAPE reagent.^37, 38^ DNA template (IDT) was amplified using Q5® High-Fidelity DNA Polymerase (NEB) following the manufacturer’s instructions using forward and reverse primer (**Table S3**). HiScribe™ T7 High Yield RNA Synthesis Kit (NEB) was used for in vitro transcription, following the manufacturer’s guidelines. RNA was purified using RNA Clean and Concentrator−5 kit (Zymo). RNA was quantified on a bioanalyzer (Agilent-RNA 6000 Nano Kit). 10 pM of the purified RNA in 12 μL of nuclease-free water was denatured at 95 for 2 min and snap-chilled. U4U6 RNA was folded using 3.3X U4U6 folding buffer (37.125 mM KPO_4_, pH 7.0, 1023 mM KCl, 16.5 mM EDTA) by annealing at 90 for 5 min before snap cooling on ice. The folded RNA was then incubated with DMSO (control), and 100 μM of compounds **1** and **22** for 20 min at RT. Each RNA was divided into two conditions, (a) DMSO treated (-) and (b) 100 mM 2A3 treated (+) and incubated at 37 for 20 min. 1 M DTT was used for quenching the reaction and RNA was column-purified using Illustra G-25 columns. Next, 1 μL of dNTPs (10 mM each, NEB) and 1 μL RT oligo (20 μM) (**Table S3**) were added to the RNA and incubated at 70 for 5 min and snap cooled in ice. 4 μL first strand buffer mix (250 mM Tris-HCl pH 8.0, 375 mM KCl), 2 μL DTT (100 mM), 1 µL RNase Inhibitor, 1 µL SuperScript II (Invitrogen), 1 µL MgCl_2_ (120 mM) were added for reverse transcription of the RNA by incubation at 25 for 5 min, followed by 42 for 2 hours and 75 for 20 min. The reverse transcribed products were purified using Illustra G-25 spin columns and used for library preparation using PCR amplification (NEB Q5® High-Fidelity DNA Polymerase and primers are listed in Table S3). The libraries were purified using DNA clean and concentrator-5 (Zymo). Illumina Miniseq instrument was used for paired-end sequencing of the libraries following the manufacturer’s instructions. The resulting fastq files were analyzed using SHAPEmapper software (https://github.com/Weeks-UNC/shapemapper2/tree/master).^39^ RNA secondary structure was predicted utilizing the SHAPE constraints using structure editor (https://rna.urmc.rochester.edu/GUI/html/StructureEditor.html). The output was also used to perform delta SHAPE analysis (https://github.com/Weeks-UNC/deltaSHAPE). Arc plots were generated using (https://github.com/Weeks-UNC/Superfold).

### Molecular Dynamics (MD) Simulations

MD simulations utilized the Amber 20/22 software package.^40^ The RNA-specific LJbb force field was used,^41^ combining the ff99OL3^42^ parameter set with the Steinbrecher and Case phosphate oxygen van der Waals radii.^43^ In addition, MD simulations employed the OPC model waters.^44, 45^ Explicit solvent molecular particle mesh Ewald (PME) dynamics simulations were utilized.^46^ In the exploratory phase multiple NMR conformers of the U4/U6 di-snRNA from the PDB: 2N7M file were used as inputs to the Amber LEaP module which combined them with waters and monovalent ions (Na+/Cl-) in cuboid solvent boxes to generate the topology and coordinate files. The minimum distance between the solute (RNA alone or RNA with the docked compounds) and solvent box boundary was set at 12 Å. The net solute charge was neutralized with Na+ ions, and additional Na+/Cl-ion pairs were added to simulate the net 0.15 M salt concentration for the entire system. Simulations were run with 2 fs time steps, employing the SHAKE algorithm to constrain all hydrogen bonds in the system. The Berendsen thermostat and algorithm were used to maintain the simulation temperature of 298 K and to maintain the pressure at 1.0 Pa in NPT simulations during system equilibration and production runs.^47^ A cut-off of 9 Å for the non-bonded interactions was used and explicit solvent periodic boundary conditions were employed.

A 12-step equilibration protocol was used that started with energy minimization of the solvent and the solute restrained, followed by multiple short phases of heating to and dynamics at 298 K, and full system energy minimizations with gradually decreasing harmonic restraints applied to the solute. The last phase of the equilibration protocol was unrestrained heating to 298 K, ramped up over 0.2 ns, and kept at the steady target temperature for 2 ns. Following equilibration, unrestrained (production) MD simulations were performed for 2 μs. Analyses of the generated MD trajectories (RMSD, and the RNA base pairing statistics) were performed with the Amber CPPTRAJ module applied to the MD production trajectories with the initial conformations used as the reference structures. The 2 μs-long trajectories were sampled in 0.1 ns steps (20,000 data points) and 1 ns step (2,000 data points) yielding statistically indistinguishable results. The Amber CPPTRAJ module was also used in the analyses of RMSD, conformation clustering and base pairing. 3D visualizations were done with the aid of PyMOL (Schrödinger, LLC. PyMOL Molecular Graphics System, Version 2.4.0. https://www.pymol.org ).

### Electrophoretic Gel Mobility Shift Assay (EMSA)

The 5′-end Cy5 labeled U4U6 RNA in 11.25 mM KPO_4_ pH 7.0, 310 mM KCl, and 5 mM EDTA was folded based on the above-mentioned method. The folded RNA was incubated with 1 µM Snu13 protein in the buffer containing 20 mM Tris-Cl pH 7.6, 150 mM NaCl, and 0.1% of Triton X-100 and 2 mM DTT for 30 min at room temperature. Then the compound (0.5 µM to 20 µM) was added to the samples and incubated at room temperature for 15 min. The samples were loaded into the 8% native PAGE gel and visualized by AMERSHAM TYPHOON Biomolecular Imager. Image J software was used for image processing. RNA-protein complex formation in the presence of increasing concentration of compound **22** was calculated by quantifying the higher molecular weight band intensity and normalized to the band intensity of RNA-protein complex in the absence of compound. Values reported are the average of three replicates. The half-maximal inhibitory concentration (IC_50_) was determined by fitting the data in inhibitor dose−response curves in GraphPad Prism 9. Quantitative data were presented as the mean ± SEM.

## RESULTS AND DISCUSSION

### Pocket identification and ligand discovery by SMM

The structure of the yeast U4/U6 snRNA assembly has previously been studied by NMR-SAXS/WAXS (PDB: 2N7M), illustrating a complex three dimensional structure ensemble.^23^ To investigate the ligandability of this complex RNA structure, we utilized Molsoft Pocket Finder (Molsoft ICM) to identify the hydrophobic regions that are potentially suitable for small molecule binding.^24, 48^ As shown in **Figure 1A** and **Table S1**, a total of 10 hydrophobic regions were identified based on the calculation. Among them, two pocket-like regions were regarded as “ligandable”, or having properties likely suitable for small molecule binding: one is in the central part of the three-way junction (3WJ, yellow) and the other is in the 5×2 asymmetric internal loop (IL), or K-turn, region (*red*) formed within the 5′ SL of U4 snRNA. The volume and drug-likeness parameters for the two pockets fell into an appropriate range (dark blue area), indicating potential for ligandability. Thus, we decided to focus on those two pockets for ligand discovery.

**Figure 1.**
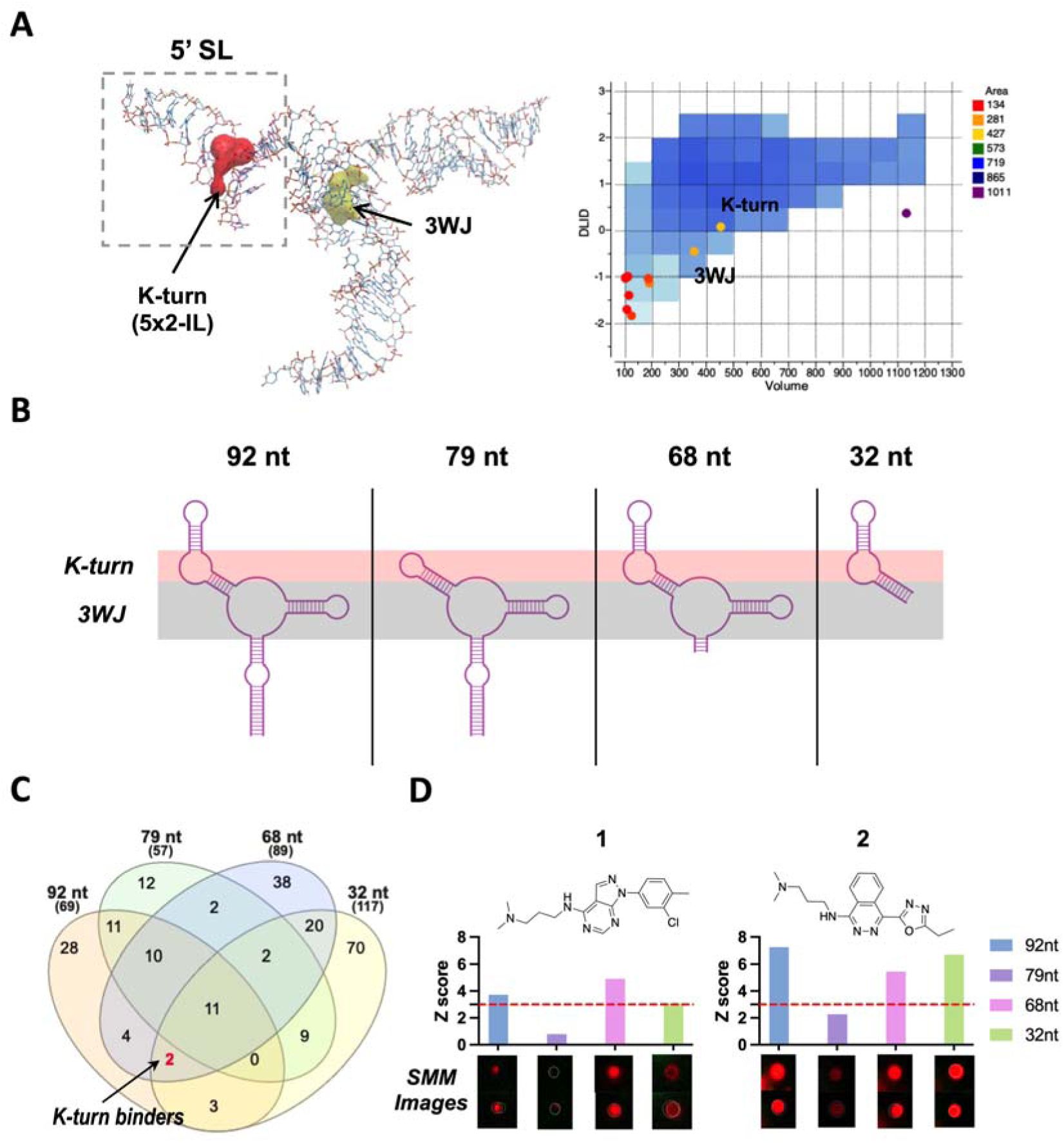
Discovery of small molecules that bind to the 5′ SL IL region in the yeast U4/U6 snRNA assembly model. (A) NMR solution structure of U4/U6 snRNA assembly mimic and hydrophobic pocket analysis (PDB: 2N7M). (B) Design of U4/U6 RNA constructs, including WT/mut/truncated RNAs for parallel SMM screening. (C) Venn diagram demonstrating SMM screening results using four different RNA constructs. (D) Chemical structure of two IL-binding molecules (**1** and **2**), as well as their corresponding Z-scores and SMM images. Red dotted line indicates hit cutoff of 3.

To target the specific regions of the U4/U6 snRNA assembly, we utilized a model of the yeast U4/U6 di-snRNA, a 92 nt RNA construct that was previously reported.^23^ We also rationally designed three additional RNA constructs that altered specific structural motifs (79 nt, 68 nt, and 32 nt) by mutating/truncating the IL or 3WJ (**Figure 1B** and **Table S2**). All four RNA samples were purchased fluorescently labeled at the 5′ end with Cy5, and used in high-throughput SMM screens in parallel as previously reported.^35, 49^ After RNA incubation, washing, and fluorescence imaging of the SMMs, Z-scores for every compound were directly compared, and theoretically, the binding specificity/binding site could be deconvoluted by the Z-score patterns, based on Z-score=3 as a cutoff. Our hypothesis was that by comparing the results of screens with closely related structures we could identify small molecules that bound to specific regions of a more complex RNA but not mutated variants. In **Figure 1C**, screening results are summarized in a Venn diagram, highlighting two compounds (**1** and **2**) that exhibited good binding selectivity in the compiled SMM dataset (**Figure S1**). According to the binding pattern (**Figure 1D**), they showed positive binding signals for K-turn-containing RNAs (92 nt, 68 nt, and 32 nt) but exhibited negative results for 79 nt RNA (lacking the K-turn), which suggested that these compounds might bind to the U4 IL-containing region. In addition, ten other compounds in the library were found to bind the 3WJ (bind 92 nt, 79 nt, and 68 nt, but not 32 nt) (**Figure S2**). However, these ligands were not selective for U4/U6 over other RNAs according to the larger compiled SMM dataset.^49^ Due to the desirable structural properties of the U4 IL, coupled with its importance in protein binding and spliceosome assembly,^23^ we decided prioritized these compounds for further study.

### Hit identification and RNA thermostability study

We utilized surface plasmon resonance (SPR) to validate the interaction between small molecules and the target RNA. Compounds were titrated to SPR surfaces containing folded U4/U6 snRNA construct (92 nt), and dose-response values were observed, resulting in K_D_ values of 50.3 ± 1.3 µM for compound **1** and 30.5 ± 8.4 µM for compound **2**, respectively (**Figure S3**). Having confirmed that the two hits could bind the RNA target, we next explored their effect on RNA stability. A thermal melting assay was performed using circular dichroism (CD). First, a CD spectrum for folded U4/U6 92 nt RNA was obtained at room temperature. The spectrum showed a strong positive peak at ∼260 nm, a minor and negative band at ∼240 nm, and an obvious negative peak at 210 nm, demonstrating a characteristic dsRNA structure. Then, a melting experiment was conducted by heating the RNA sample from 20 to 90 . As shown in **Figure 2A**, during melting, the peaks at 260 nm and 210 nm both decreased, while the band at 240 nm became much broader. This change is attributed to the unwinding of the right-handed helix structure and the disruption of base pairing.^50^ Moreover, the 260 nm peak also gradually shifted towards a higher wavelength, ending up at ∼270 nm, consistent with single-stranded RNA. By tracking the intensity of the 260 nm peak, a melting curve can be obtained, and the melting temperature (T_m_) for an RNA construct can be calculated based on the first derivative (**Figure 2B and 2C**). As a result, the T_m_ for the 92 nt RNA was found to be 65.6 ± 0.3 . Upon addition of an excess (100 µM) of **1**, the T_m_ slightly decreased (ΔT_m_ = -1.5 ± 0.5 ), while adding the same amount of **2** resulted in stabilization of the U4/U6 RNA (ΔT_m_ = +3.3 ± 0.3 ). This indicated that the two compounds might have different binding modes, although they both showed binding preference to the IL and had modest but reproducible effects on unfolding temperature. Given that there are multiple structural features of this RNA it is likely a complex unfolding event, and more in-depth interpretation of the melting with CD is likely to be difficult.

**Figure 2.**
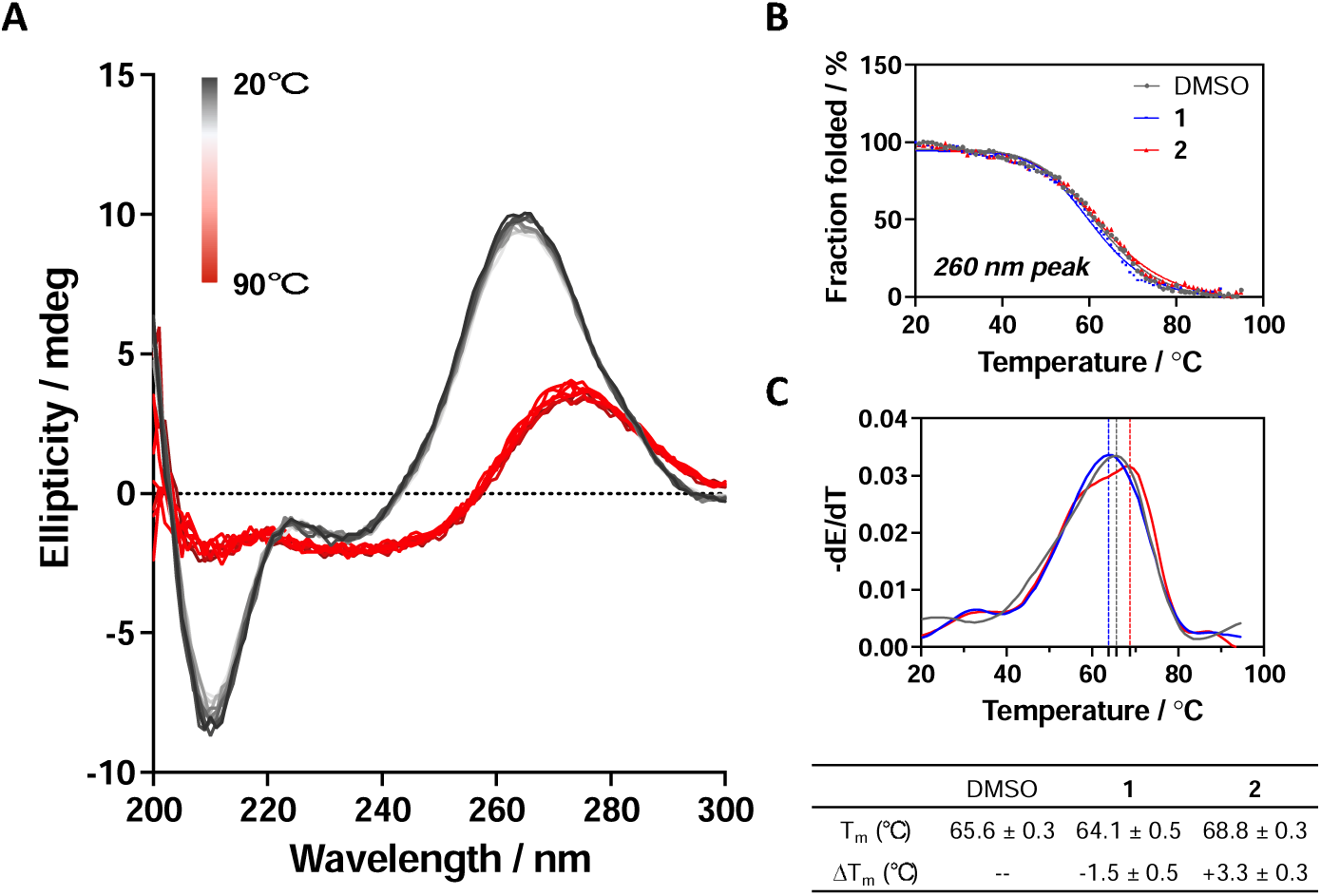
RNA stability study in the presence/absence of small molecules. (A) Temperature-dependent CD spectra of the U4/U6 snRNA model. (B) CD melting curves by tracing the peaks at 260 nm with/without adding 100 µM compound. (C) T_m_ determination using the first derivatives of the CD melting curves.

### SAR study and Lead optimization

We further performed structure-activity relationship (SAR) studies on a focused compound library to improve the probes’ binding affinity. We started with analogs of compound **2** since it was a slightly tighter binder. Compounds similar to **2** were purchased or synthesized, and the binding between analogs and U4/U6 snRNA was measured by SPR. As shown in **Table 1** and **Figure S4**, the compounds exhibit a variety of K_D_ values from low micromolar interaction to no binding, revealing a clear SAR. Among the newly tested 26 analogs, compound **22** showed the tightest binding to the 92 nt RNA construct, with a K_D_ of 3.9 ± 2.2 µM (**Figure 3A and 3B**). More importantly, when using 79 nt RNA that disrupts the 5′ SL, this compound did not show any binding until 50 µM (**Figure S5**), indicating a good binding selectivity towards the IL region. Meanwhile, the binding between **22** and U4/U6 snRNA was also tested by MST as a cross-validation, resulting in a K_D_ of 9.1 ± 0.6 µM (**Figure 3C**), which is in good agreement with the SPR data.

**Figure 3.**
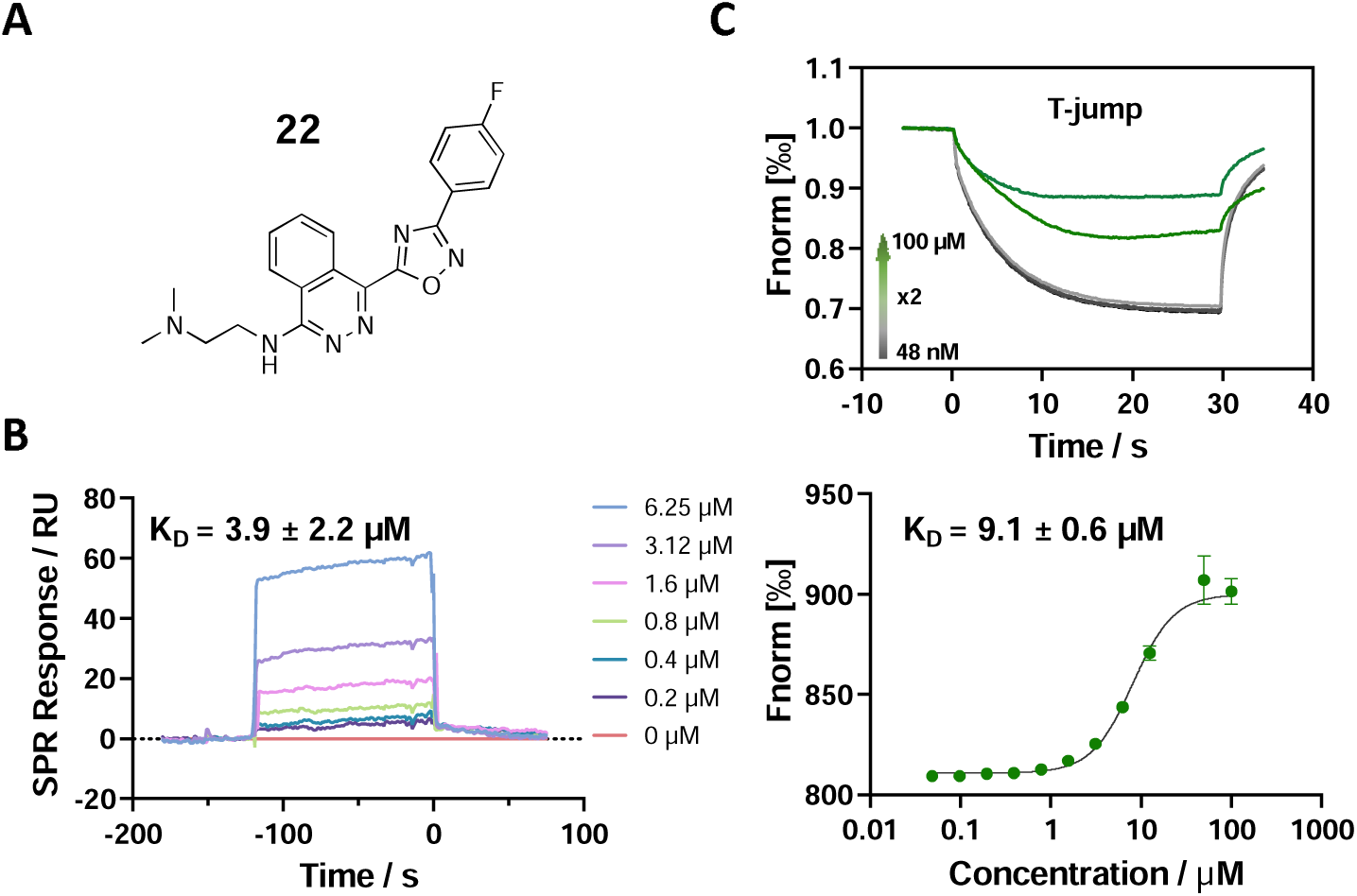
Binding affinity measurement of the optimized IL-binding compound. (A) chemical structure of **22**. (B) SPR binding curves of compound titration on RNA-coated chip. (C) MST trace and dose-dependent response of compound **22** binding with U4/U6 92-nt RNA construct. F_norm_ data are presented as mean ± SD (n=3).

**Table 1.** Structure-activity relationship (SAR) study on SMM hit 2.

| ID | R<sub>1</sub> | Het | R<sub>2</sub> | SPR K<sub>D</sub><br>($\mu\text{M}$)<sup>a</sup> | ID | R<sub>1</sub> | Het | R<sub>2</sub> | SPR K<sub>D</sub><br>($\mu\text{M}$)<sup>a</sup> |
| --- | --- | --- | --- | --- | --- | --- | --- | --- | --- |
| **2<sup>b</sup>** | | | | $30.5 \pm 8.4$ | **14** | | | | $32.8 \pm 11.0$ |
| **3** | | | | $29.2 \pm 8.6$ | **15** | | | | $23.4 \pm 6.0$ |
| **4** |  |  |  | >100 | **16** |  |  |  | N/A |
| **5** |  |  |  | N/A | **17** |  |  |  | N/A |
| **6** |  |  |  | N/A | **18** |  |  |  | N/A |
| **7** | | | | N/A | **19** | | | | $34.5 \pm 11.3$ |
| **8** | | | | N/A | **20** | | | | $42.1 \pm 8.6$ |
| **9** | | | | $61.3 \pm 38.9$ | **21** | | | | $23.0 \pm 5.3$ |
| **10** | | | | $51.4 \pm 8.0$ | **22** | | | | $3.9 \pm 2.2$ |
| **11** | | | | N/A | **23** | | | | $11.9 \pm 7.1$ |
| **12** | | | | $24.2 \pm 7.0$ | **24** | | | | >100 |
| **13** | | | | N/A | **25** | | | | $13.9 \pm 3.2$ |

- a. For $K_D$ values determined by SPR, N/A suggests that there was no binding observed during the whole titration (0~100 $\mu\text{M}$ ); meanwhile, $K_D > 100 \mu\text{M}$ means that the binding signal appeared at 100 $\mu\text{M}$ but failed to fit the curves. - b. Original hit from SMM screening.

### Structure probing by SHAPE-MaP

To investigate the conformational effects of small molecules on U4/U6 RNA structure, we performed structural probing experiments using SHAPE-MaP.^51^ This approach reports on the flexibility of each nucleotide within the U4/U6 snRNA structure and how it changes upon binding and interaction with **22**. Chemical probing using 2A3 coupled with high-throughput sequencing was performed. The SHAPE reactivity of experimentally derived nucleotide constraints was obtained using SHAPEmapper^39^ and used to predict the base pairing interactions of the RNA and the effect of each compound on the secondary structure. The results of RNA structure prediction (MFE and centroid) were visualized by the RNAfold Webserver using SHAPE reactivity as an additional restraint (http://rna.tbi.univie.ac.at/cgi-bin/RNAWebSuite/RNAfold.cgi). Compounds **1** and **22** showed the opposite effects on U4U6 RNA folding when compared to DMSO (**Figure S6**). According to base pairing probabilities derived from SHAPE reactivity, **1** exhibited a clear destabilization effect on the 5′ SL as well as the Stem 1 region. Distinct from compound **1**, **22** demonstrated an overall stabilization of the RNA structure, centered around bases within the K-turn region in 5′ SL. These observations agreed with the CD melting data reported above. Neither compound resulted in the rearrangement of predicted base pairing interactions of the RNA.

Next, deltaSHAPE was utilized to analyze detailed effects of RNA-binding small molecules. Specifically, in when bound with **22**, the U4/U6 RNA showed enhanced stability in the 5′ SL region when compared to the unbound state (**Figure 4A**). This could thus plausibly be the binding region for **22**. Arc plots derived from the SHAPE-MaP data depict the base-pairing probability changes in the presence and absence of **22** (**Figure 4B**). DeltaSHAPE results revealed two regions that are significantly altered upon compound binding, specifically C28, A29, and G30 in the 5′ SL and G13, G15, and A16 belonging to the Stem 2 domain (**Figure 4C**). The former region affected by **22** is in the K-turn motif as expected. The latter one is near 3WJ, which might be due to the long-range effect of the conformational change in 5′ SL, potentially resulting in slight rearrangement of basepairing near the 3WJ.

**Figure 4.**
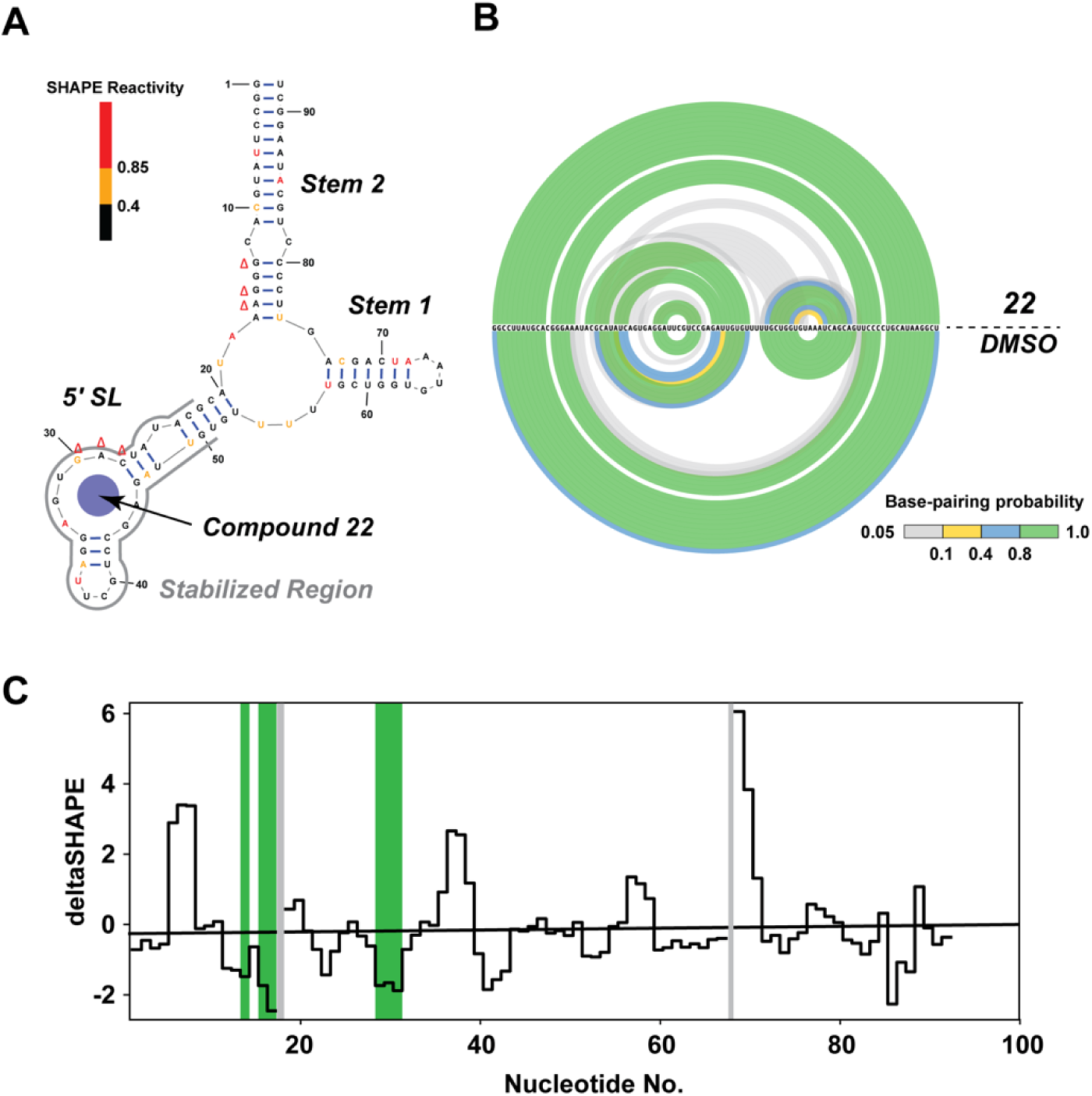
Effects of compound **22** on the secondary structure of U4/U6 snRNA. (A) Secondary structure of the U4/U6 snRNA construct. Red, yellow, and black bars represent high, moderate, and low SHAPE reactivity, while red “Δ” symbols suggest observed change at individual nucleotides. (B) Arc plots for the comparison of base-pairing probabilities between compound- and DMSO-treated conditions. Base-pairing probability is depicted in the color scheme shown at the bottom. (C) Skyline plots visualizing the delta SHAPE data. The highlighted regions in green show significant changes in corresponding positions. The highlighted grey regions represent nucleotides with low SHAPE reactivity.

### MD simulation analysis on RNA-small molecule interaction

To further study the stabilization effect of compound **22** on the RNA, MD simulations were performed utilizing the reported structural information of U4/U6 snRNA (PDB: 2N7M, solution NMR). First, the compound was docked into the 5×2-IL (K-turn region) using NMR frame 1 conformation as the starting state, resulting in a target-stabilizing conformation with a score of -15.96 kcal/mol. An MD simulation was then launched based on the initial docked pose. At the beginning of the simulation, compound **22** immediately started to adjust, intercalating itself into the loop, passing through two relatively intermediate states, and stabilizing past 1,080 ns (**Figure S7**). As shown in **Figure 5A**, the phthalazine core of the compound is stacked with A45, while the benzyl ring interacts with G32 (parallel) and G30 (perpendicular). With time passing, the loop becomes twisted due to the adjustment, and this final pose is stable from 1,080 ns to 2,000 ns. To quantitatively analyze, the 5′ SL RNA was divided into smaller sections, including duplex A (20-24:49-53), duplex B (26-28:46-48), 5×2-IL (29-33:44-45), duplex C (34-36:41-43), and hairpin H (37-40) (**Figure 5B**). RMSD data were calculated for all heavy atoms in each section and compared. For the *apo* RNA, the RMSD of the 5×2-IL region was 5.3 Å, significantly higher than the levels in sections A (2.4 Å), B (2.2 Å), and C (2.1 Å), indicating that the loop has more flexibility than the three duplexes. With compound bound, the RMSD of the loop greatly decreased to 3.0 Å while the three duplexes remained stable with low RMSD (2.1∼2.3 Å). This suggested that the compound interacted with the loop region and stabilized the structure. In addition, by analyzing the base-pairing frequency of sections A and B, it is worth noting that three base pairs (20A-53U, 24A-49U, and 26A-48U) were strengthened (**Figure 5C and S8**). This suggested that the stabilization of the internal loop by the small molecule could potentially have a long-range effect on the region near 3WJ, thus globally impacting RNA folding. This observation is also consistent with the SHAPE reactivity result (**Figure S6**). Overall, the results of the MD simulation are consistent with the SHAPE-MaP result, demonstrating a stabilization effect of compound **22** on the RNA. Thus, the MD analysis provides further insights into how the molecule interacts with the internal loop and influences the stability of the base pairing near the loop.

**Figure 5.**
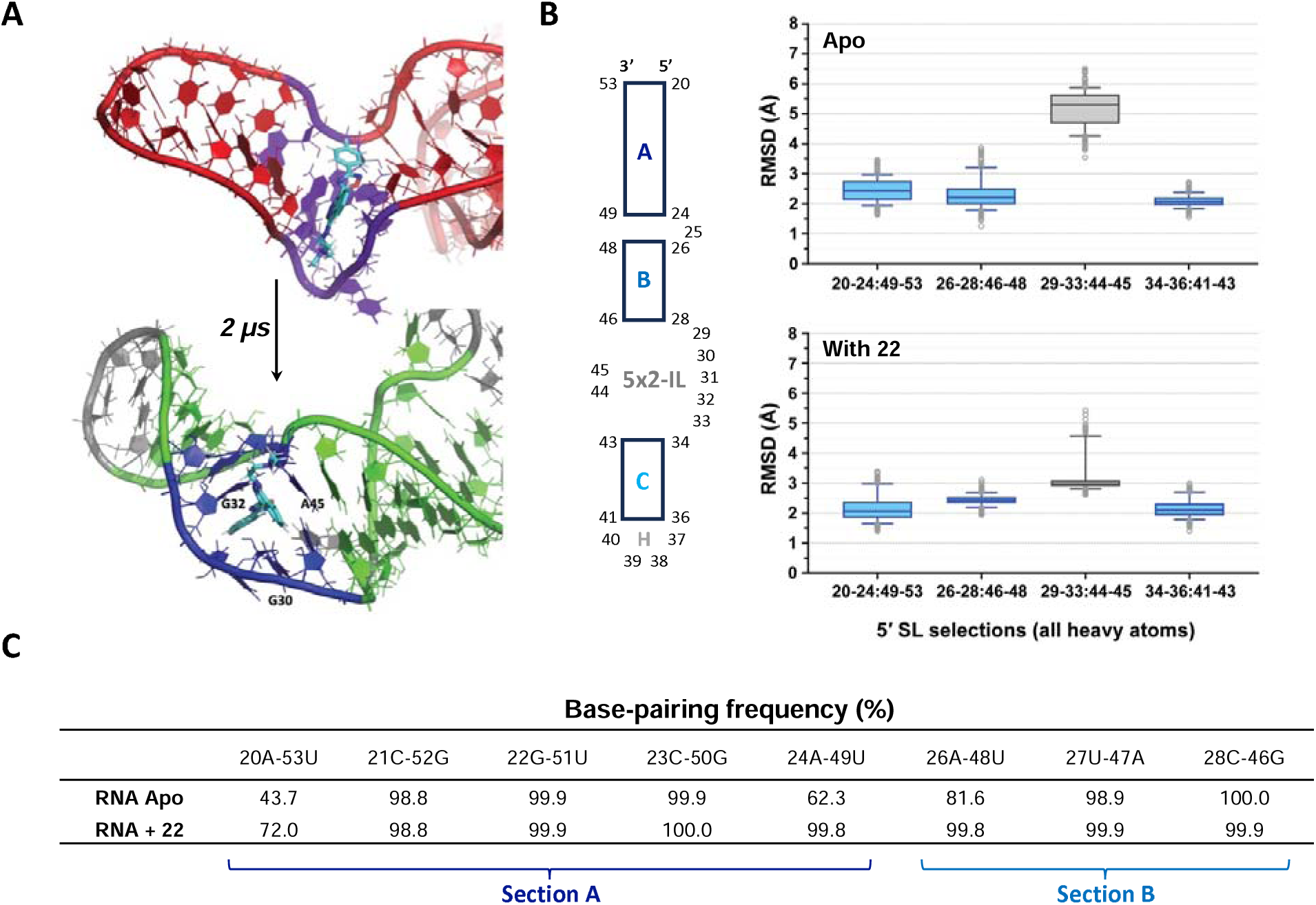
MD simulation analysis on 5′ SL interacting with compound **22**. (A) Docking **22** into U4 snRNA 5×2-IL (K-turn, PDB: 2N7M). MD simulation demonstrating the initial pose (top) and final stable pose at 2,000 ns (bottom). (B) The RMSD data for segments (A, B, L, C) in the 5′ SL in the absence or presence of the compound. (C) Base-pairing frequency analysis on the RNA segments without/with the compound.

**Figure 6.**
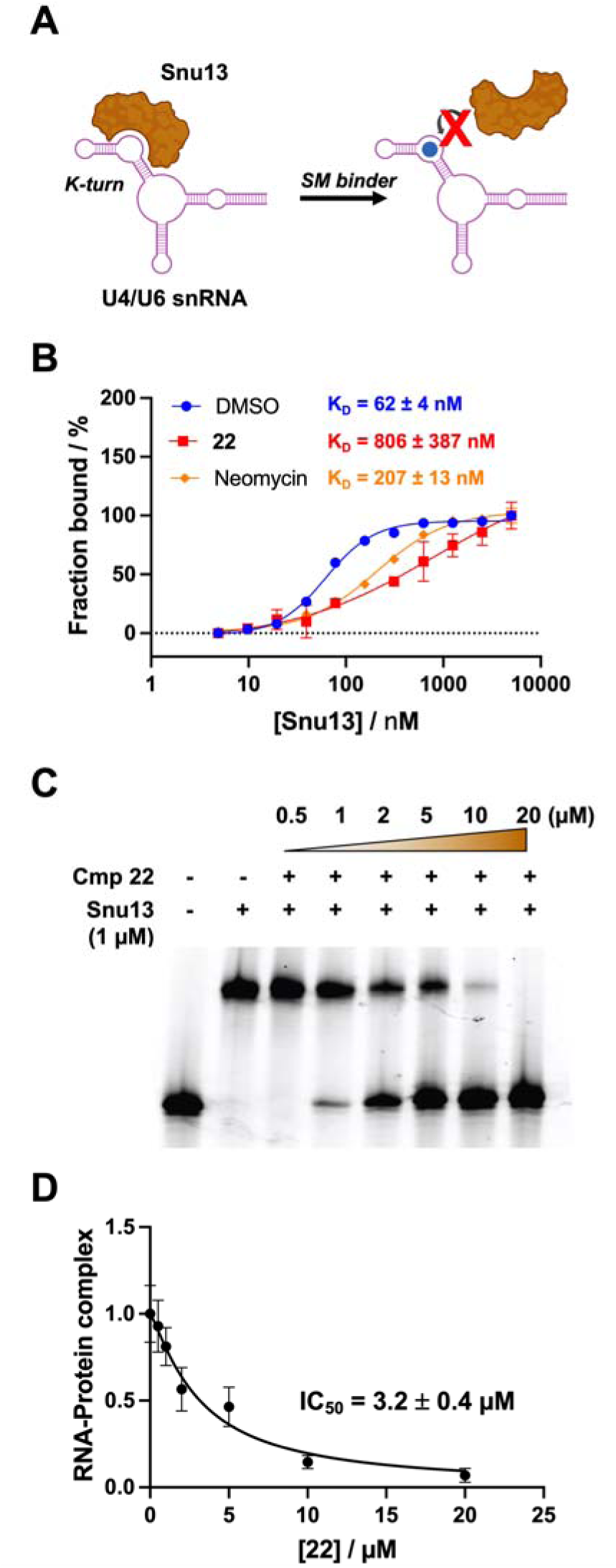
(A) *In vitro* inhibition of U4/U6 snRNA-Snu13 binding by compound **22**. (B) K_D_ shift assay by MST suggests the inhibition of protein-RNA interaction. 50 nM of 5′-Cy5-labeled U4/U6 RNA construct (92 nt) was tested. 10 µM of **22** or neomycin was added to the solution during MST titration. (C) EMSA results demonstrate concentration-dependent inhibition of RNA-protein interaction with the addition of compound **22**. (D) IC_50_ determination by quantification of gel band intensity. Error bars represent the standard deviation from three independent replicates.

### Inhibitory effect of 22 on RNA-protein interaction

At the precatalytic stage of RNA splicing, U4/U6.U5 tri-snRNP is assembled, an essential step in the formation of the B complex in the spliceosome. The structure of yeast U4/U6.U5 tri-snRNP solved by Cryo-electron microscopy (Cryo-EM) provides atomic resolution information about the U4/U6 di-snRNA and its interaction with several core proteins including Snu13, Prp3, Prp4, and Prp31.^52^ Among them, Snu13 recognizes the 5′ SL of U4 snRNA and binds to the K-turn region, resulting in a conformational change and modulating the dynamics of U4/U6 di-snRNP.^53^ This observation is confirmed by the structural comparison between free U4/U6 di-snRNA and the corresponding snRNP complex. In brief, this conformational change of RNA at K-turn caused by protein binding leads to the recruitment of Prp31 to 5′ SL, followed by the recruitment of Prp3 and Prp4, which then remodels the U4/U6 interhelix junction.

We tested whether the binding of the compound had any effect on the interaction between the U4/U6 model RNA and Snu13. We first performed a K_D_ shift assay by microscale thermophoresis (MST). In the absence of compound **22**, the RNA-protein interaction was characterized by titrating Snu13 into 50 nM Cy5-labeled U4/U6 RNA, reporting an affinity of 62 ± 4 nM. As a control, we used neomycin, a non-specific RNA IL-binding molecule, also a reported yeast pre-mRNA splicing inhibitor.^54^ In the presence of 10 µM neomycin, the K_D_ of RNA-Snu13 interaction was measured to be 207 ± 13 nM, demonstrating that small molecules are capable of altering the binding of Snu13. In the presence of 10 µM **22** in the working solution, the binding affinity of Snu13 was also shifted, resulting in an affinity of 806 ± 387 nM, 13-fold weaker. This result suggested that **22** competes with Snu13 and alters binding affinity between the protein and RNA U4 IL. IL

To further quantify the effect of compound **22** on the Snu13-U4/U6 RNA interaction, we conducted an EMSA experiment. Knowing the approximate binding affinity of the Snu13-U4/U6 interaction, we used 1 µM of protein to saturate the binding with Cy5-labeled RNA. Then, solutions of **22** were added at different concentrations ranging from 0.5 µM to 20 µM. In the absence of **22**, only the RNA-protein complex (upper band) was observed. As a function of the concentration of **22**, the upper band corresponding to the complex gradually faded, while the lower band demonstrating free RNA became increasingly intense. This result again confirmed the inhibition effect on the protein recruitment, caused by the binding of the compound. An IC_50_ of 3.2 ± 0.4 µM was obtained, consistent with the K_D_ value determined by SPR binding assay.

Binding assays performed by two orthogonal techniques suggested that ligand **22** could be potentially utilized as a probe to interfere with Snu13-U4/U6 interaction, thus modulating the U4/U6.U5 tri-snRNP assembly, a key step in spliceosome assembly that occurs immediately prior to catalysis. Moreover, by aligning and comparing the RNA sequences between yeast and human snRNAs, there is over 70% sequence similarity in the 5′ SL region (**Figure S9**), and the protein-binding conformations of the IL are quite similar according to Cryo-EM structures (**Figure S10**).^52, 55^ Thus, this probe also has the potential to be used as a probe for the human spliceosome. Previously, topotecan was discovered to inhibit the interaction between NHP2L1 and U4 5′ SL by RNA binding.^29^ However, the binding affinity was not determined due to the difficulty of detection by SPR and topotecan is known to broadly interact with diverse nucleic acid structures.^56, 57^ The discovery of compound **22** provides us with an alternative tool to modulate spliceosome assembly and understand RNA-protein interactions within the spliceosome, and more studies on in-cell functions (such as alternative splicing and protein expression) will be pursued in the future.

## CONCLUSIONS

In recent years, RNA as a target for small molecules has become an intriguing topic in the field of drug discovery. Different approaches of high-throughput screening have been established for RNA targeting, such as MS (e.g., ALIS developed by Merck & Co., Inc.),^58^ NMR,^59, 60^, FRET,^61^ SPR,^62^ and multiple SMM-based approaches including 2-dimensional combinatorial screening and INFORNA.^33, 63–65^ To avoid non-specific binders, multiple RNA targets or RNA complexes (e.g., tRNA) are used as controls, and criteria for specificity (e.g., Gini coefficient) have been utilized.^66^ However, most approaches provide target-specific information, but not structure-specific information for where a hit ligand binds within a complex RNA. This study provides a novel strategy for screening related constructs with mutations in specific structural regions on SMMs, to target a particular region of a complex RNA structure. Based on this screening approach, two compounds showed specific binding to the 5′ SL structure in U4/U6 snRNA assembly. Interestingly, they exhibited divergent effects on RNA thermodynamic stability as characterized by CD melting assay (**1**, destabilization; **2**, stabilization). This observation indicated that there might be two distinct binding modes of the two molecules, was later confirmed by SHAPE-MaP experiments. By focusing on an SAR study of compound **2**, one of the analogues (**22**) exhibited a greatly improved K_D_ of RNA binding. DeltaSHAPE analysis revealed that the compound bound the K-turn region in U4 5′ SL, due to the change of SHAPE reactivity in that region (especially C28, A29, and G30). MD simulations using the published U4/U6 RNA structure solved by NMR-SAXS/WAXS (PDB: 2N7M) also aligned with the experimental results. Moreover, in the presence of **22**, the IL of the RNA exhibited a distinct conformation from the *apo* state and protein-binding state, indicating that the ligand binding could affect the downstream protein recruitment by preventing the formation of the protein-binding conformation. This hypothesis was validated by both the K_D_-shift assay by MST and the competitive assay by EMSA. As a result, the compound could inhibit the binding of Snu13.

Overall, this study demonstrates the discovery of multiple new small molecule ligands for U4/U6 snRNA targeting to modulate RNA-protein interactions that are important steps in spliceosome assembly. We identify multiple distinct small molecules with divergent effects on the flexibility of the same binding site, highlighting a complexity of using binding assays as a primary screen. SHAPE-MaP, paired with molecular dynamics simulations, revealed insights into how these chemical classes impact conformational dynamics of the RNA. Here, the hit compound that resulted in an overall thermodynamic stabilization of the RNA was successfully optimized into an inhibitor of an RNA-protein interaction *in vitro*. More broadly, the high-throughput screening approach described here will likely be valuable for future studies aimed toward significant to rapid development of site- and pocket-specific RNA-binding molecules. While the compounds reported here may be insufficiently potent to impact splicing *in vivo*, our studies reveal that an approach targeting a specific hydrophobic pocket within an RNA is a valid strategy to identify small molecules capable of disrupting specific RNA-protein interactions. Finally, beyond this target, the approach of counterscreening mutant/wild type constructs to rapidly identify site-specific ligands will likely have broader applications in the RNA-targeting field as increasingly complex RNAs are considered as targets.

## ASSOCIATED CONTENT

### Supporting Information

The Supporting Information is available free of charge on the xxx website at DOI: xxxxxx

## AUTHOR INFORMATION

### Notes

The authors declare no competing financial interest.

## ACKNOWLEDGMENTS

This research was supported by the Intramural Research Program of the National Institutes of Health, National Cancer Institute (NCI), Center for Cancer Research (1ZIABC011585). The authors thank the members of the biophysics resource (Dr. Sergey G. Tarasov and Marzena Dyba) for helpful comments and suggestions on biophysical experiments. We thank Dr. Joel P. Schneider from NCI/CBL for providing access to the SPR instrument. We thank Dr. Patrick S. Irving from Prof. Kevin M. Weeks’ lab at UNC for useful discussion on SHAPE data presentation. This work utilized the computational resources of the NIH HPC Biowulf cluster (http://hpc.nih.gov).

This project has been funded in whole or in part with Federal funds from the National Cancer Institute, National Institutes of Health, Department of Health and Human Services, under Contract No. 75N91019D00024 and 75N91024F00011. The content of this publication does not necessarily reflect the views or policies of the Department of Health and Human Services, nor does mention of trade names, commercial products, or organizations imply endorsement by the U.S. Government.

## REFERENCES

(1) Wahl, M. C.; Will, C. L.; Luhrmann, R. The spliceosome: design principles of a dynamic RNP machine. Cell 2009, 136 (4), 701–718. DOI: 10.1016/j.cell.2009.02.009 From NLM Medline.

(2) Choquet, K.; Patop, I. L.; Churchman, L. S. The regulation and function of post-transcriptional RNA splicing. Nat Rev Genet 2025, 26 (6), 378–394. DOI: 10.1038/s41576-025-00836-z From NLM Medline.

(3) Wright, C. J.; Smith, C. W. J.; Jiggins, C. D. Alternative splicing as a source of phenotypic diversity. Nat Rev Genet 2022, 23 (11), 697–710. DOI: 10.1038/s41576-022-00514-4 From NLM Medline.

(4) Bradley, R. K.; Anczukow, O. RNA splicing dysregulation and the hallmarks of cancer. Nat Rev Cancer 2023, 23 (3), 135–155. DOI: 10.1038/s41568-022-00541-7 From NLM Medline.

(5) Barraza, S. J.; Bhattacharyya, A.; Trotta, C. R.; Woll, M. G. Targeting strategies for modulating pre-mRNA splicing with small molecules: Recent advances. Drug Discov Today 2023, 28 (1), 103431. DOI: 10.1016/j.drudis.2022.103431 From NLM Medline.

(6) Ghosh, A. K.; Mishevich, J. L.; Jurica, M. S. Spliceostatins and Derivatives: Chemical Syntheses and Biological Properties of Potent Splicing Inhibitors. J Nat Prod 2021, 84 (5), 1681–1706. DOI: 10.1021/acs.jnatprod.1c00100 From NLM Medline.

(7) Childs-Disney, J. L.; Yang, X.; Gibaut, Q. M. R.; Tong, Y.; Batey, R. T.; Disney, M. D. Targeting RNA structures with small molecules. Nat Rev Drug Discov 2022, 21 (10), 736–762. DOI: 10.1038/s41573-022-00521-4 From NLM Medline.

(8) Fullenkamp, C. R.; Liang, X.; Pettersson, M.; Schneekloth Jr., J. Outlook. In RNA as a Drug Target, 2024; pp 355–384.

(9) Falese, J. P.; Donlic, A.; Hargrove, A. E. Targeting RNA with small molecules: from fundamental principles towards the clinic. Chem Soc Rev 2021, 50 (4), 2224–2243. DOI: 10.1039/d0cs01261k From NLM Medline.

(10) Zhang, Z.; Pinto, A. M.; Wan, L.; Wang, W.; Berg, M. G.; Oliva, I.; Singh, L. N.; Dengler, C.; Wei, Z.; Dreyfuss, G. Dysregulation of synaptogenesis genes antecedes motor neuron pathology in spinal muscular atrophy. Proc Natl Acad Sci U S A 2013, 110 (48), 19348–19353. DOI: 10.1073/pnas.1319280110 From NLM Medline.

(11) Neueder, A.; Dumas, A. A.; Benjamin, A. C.; Bates, G. P. Regulatory mechanisms of incomplete huntingtin mRNA splicing. Nat Commun 2018, 9 (1), 3955. DOI: 10.1038/s41467-018-06281-3 From NLM Medline.

(12) Michaels, W. E.; Pena-Rasgado, C.; Kotaria, R.; Bridges, R. J.; Hastings, M. L. Open reading frame correction using splice-switching antisense oligonucleotides for the treatment of cystic fibrosis. Proc Natl Acad Sci U S A 2022, 119 (3). DOI: 10.1073/pnas.2114886119 From NLM Medline.

(13) Dhillon, S. Risdiplam: First Approval. Drugs 2020, 80 (17), 1853–1858. DOI: 10.1007/s40265-020-01410-z.

(14) Ratni, H.; Ebeling, M.; Baird, J.; Bendels, S.; Bylund, J.; Chen, K. S.; Denk, N.; Feng, Z.; Green, L.; Guerard, M.;, et al. Discovery of Risdiplam, a Selective Survival of Motor Neuron-2 ( SMN2) Gene Splicing Modifier for the Treatment of Spinal Muscular Atrophy (SMA). J Med Chem 2018, 61 (15), 6501–6517. DOI: 10.1021/acs.jmedchem.8b00741 From NLM Medline.

(15) Saade, J.; Mestre, T. A. Huntington’s Disease: Latest Frontiers in Therapeutics. Curr Neurol Neurosci Rep 2024. DOI: 10.1007/s11910-024-01345-y From NLM Publisher.

(16) Chen, J. L.; Zhang, P.; Abe, M.; Aikawa, H.; Zhang, L.; Frank, A. J.; Zembryski, T.; Hubbs, C.; Park, H.; Withka, J.;, et al. Design, Optimization, and Study of Small Molecules That Target Tau Pre-mRNA and Affect Splicing. J Am Chem Soc 2020, 142 (19), 8706–8727. DOI: 10.1021/jacs.0c00768 From NLM Medline.

(17) Araki, S.; Ohori, M.; Yugami, M. Targeting pre-mRNA splicing in cancers: roles, inhibitors, and therapeutic opportunities. Front Oncol 2023, 13, 1152087. DOI: 10.3389/fonc.2023.1152087 From NLM PubMed-not-MEDLINE.

(18) Tang, Z.; Zhao, J.; Pearson, Z. J.; Boskovic, Z. V.; Wang, J. RNA-Targeting Splicing Modifiers: Drug Development and Screening Assays. Molecules 2021, 26 (8). DOI: 10.3390/molecules26082263 From NLM Medline.

(19) Boer, R. E.; Torrey, Z. R.; Schneekloth, J. S., Jr. Chemical Modulation of Pre-mRNA Splicing in Mammalian Systems. ACS Chem Biol 2020, 15 (4), 808–818. DOI: 10.1021/acschembio.0c00001 From NLM Medline.

(20) Effenberger, K. A.; Urabe, V. K.; Jurica, M. S. Modulating splicing with small molecular inhibitors of the spliceosome. Wiley Interdiscip Rev RNA 2017, 8 (2). DOI: 10.1002/wrna.1381 From NLM Medline.

(21) Wilkinson, M. E.; Charenton, C.; Nagai, K. RNA Splicing by the Spliceosome. Annu Rev Biochem 2020, 89, 359–388. DOI: 10.1146/annurev-biochem-091719-064225 From NLM Medline.

(22) Nguyen, T. H.; Galej, W. P.; Bai, X. C.; Savva, C. G.; Newman, A. J.; Scheres, S. H.; Nagai, K. The architecture of the spliceosomal U4/U6.U5 tri-snRNP. Nature 2015, 523 (7558), 47–52. DOI: 10.1038/nature14548 From NLM Medline.

(23) Cornilescu, G.; Didychuk, A. L.; Rodgers, M. L.; Michael, L. A.; Burke, J. E.; Montemayor, E. J.; Hoskins, A. A.; Butcher, S. E. Structural Analysis of Multi-Helical RNAs by NMR-SAXS/WAXS: Application to the U4/U6 di-snRNA. J Mol Biol 2016, 428 (5 Pt A), 777–789. DOI: 10.1016/j.jmb.2015.11.026 From NLM Medline.

(24) Hewitt, W. M.; Calabrese, D. R.; Schneekloth, J. S., Jr. Evidence for ligandable sites in structured RNA throughout the Protein Data Bank. Bioorg Med Chem 2019, 27 (11), 2253–2260. DOI: 10.1016/j.bmc.2019.04.010 From NLM Medline.

(25) Veenbaas, S. D.; Koehn, J. T.; Irving, P. S.; Lama, N. N.; Weeks, K. M. Ligand-binding pockets in RNA and where to find them. Proc Natl Acad Sci U S A 2025, 122 (17), e2422346122. DOI: 10.1073/pnas.2422346122 From NLM Medline.

(26) Falb, M.; Amata, I.; Gabel, F.; Simon, B.; Carlomagno, T. Structure of the K-turn U4 RNA: a combined NMR and SANS study. Nucleic Acids Res 2010, 38 (18), 6274–6285. DOI: 10.1093/nar/gkq380 From NLM Medline.

(27) Huang, L.; Wang, J.; Lilley, D. M. A critical base pair in k-turns determines the conformational class adopted, and correlates with biological function. Nucleic Acids Res 2016, 44 (11), 5390–5398. DOI: 10.1093/nar/gkw201 From NLM Medline.

(28) McPhee, S. A.; Huang, L.; Lilley, D. M. A critical base pair in k-turns that confers folding characteristics and correlates with biological function. Nat Commun 2014, 5, 5127. DOI: 10.1038/ncomms6127 From NLM Medline.

(29) Diouf, B.; Lin, W.; Goktug, A.; Grace, C. R. R.; Waddell, M. B.; Bao, J.; Shao, Y.; Heath, R. J.; Zheng, J. J.; Shelat, A. A.;, et al. Alteration of RNA Splicing by Small-Molecule Inhibitors of the Interaction between NHP2L1 and U4. SLAS Discov 2018, 23 (2), 164–173. DOI: 10.1177/2472555217735035 From NLM Medline.

(30) Warner, K. D.; Hajdin, C. E.; Weeks, K. M. Principles for targeting RNA with drug-like small molecules. Nat Rev Drug Discov 2018, 17 (8), 547–558. DOI: 10.1038/nrd.2018.93 From NLM Medline.

(31) Aguilar, R.; Spencer, K. B.; Kesner, B.; Rizvi, N. F.; Badmalia, M. D.; Mrozowich, T.; Mortison, J. D.; Rivera, C.; Smith, G. F.; Burchard, J.;, et al. Targeting Xist with compounds that disrupt RNA structure and X inactivation. Nature 2022, 604 (7904), 160–166. DOI: 10.1038/s41586-022-04537-z From NLM Medline.

(32) Vegas, A. J.; Fuller, J. H.; Koehler, A. N. Small-molecule microarrays as tools in ligand discovery. Chem Soc Rev 2008, 37 (7), 1385–1394. DOI: 10.1039/b703568n From NLM Medline.

(33) Abulwerdi, F. A.; Schneekloth, J. S., Jr. Microarray-based technologies for the discovery of selective, RNA-binding molecules. Methods 2016, 103, 188–195. DOI: 10.1016/j.ymeth.2016.04.022 From NLM Medline.

(34) Yang, M.; Carter, S.; Parmar, S.; Bume, D. D.; Calabrese, D. R.; Liang, X.; Yazdani, K.; Xu, M.; Liu, Z.; Thiele, C. J.;, et al. Targeting a noncanonical, hairpin-containing G-quadruplex structure from the MYCN gene. Nucleic Acids Res 2021, 49 (14), 7856–7869. DOI: 10.1093/nar/gkab594 From NLM Medline.

(35) Prestwood, P. R.; Yang, M.; Lewis, G. V.; Balaratnam, S.; Yazdani, K.; Schneekloth, J. S., Jr. Competitive Microarray Screening Reveals Functional Ligands for the DHX15 RNA G-Quadruplex. ACS Med Chem Lett 2024, 15 (6), 814–821. DOI: 10.1021/acsmedchemlett.3c00574 From NLM PubMed-not-MEDLINE.

(36) Moon, M. H.; Hilimire, T. A.; Sanders, A. M.; Schneekloth, J. S., Jr. Measuring RNA-Ligand Interactions with Microscale Thermophoresis. Biochemistry 2018, 57 (31), 4638–4643. DOI: 10.1021/acs.biochem.7b01141 From NLM Medline.

(37) Smola, M. J.; Rice, G. M.; Busan, S.; Siegfried, N. A.; Weeks, K. M. Selective 2’-hydroxyl acylation analyzed by primer extension and mutational profiling (SHAPE-MaP) for direct, versatile and accurate RNA structure analysis. Nat Protoc 2015, 10 (11), 1643–1669. DOI: 10.1038/nprot.2015.103 From NLM Medline.

(38) Incarnato, D. Sequencing-based analysis of RNA structures in living cells with 2A3 via SHAPE-MaP. Methods Enzymol 2023, 691, 153–181. DOI: 10.1016/bs.mie.2023.03.021 From NLM Medline.

(39) Busan, S.; Weeks, K. M. Accurate detection of chemical modifications in RNA by mutational profiling (MaP) with ShapeMapper 2. RNA 2018, 24 (2), 143–148. DOI: 10.1261/rna.061945.117 From NLM Medline.

(40) Case, D. A.; Cheatham, T. E., 3rd; Darden, T.; Gohlke, H.; Luo, R.; Merz, K. M., Jr.; Onufriev, A.; Simmerling, C.; Wang, B.; Woods, R. J. The Amber biomolecular simulation programs. J Comput Chem 2005, 26 (16), 1668–1688. DOI: 10.1002/jcc.20290 From NLM Medline.

(41) Bergonzo, C.; Cheatham, T. E., 3rd. Improved Force Field Parameters Lead to a Better Description of RNA Structure. J Chem Theory Comput 2015, 11 (9), 3969–3972. DOI: 10.1021/acs.jctc.5b00444 From NLM Medline.

(42) Zgarbova, M.; Otyepka, M.; Sponer, J.; Mladek, A.; Banas, P.; Cheatham, T. E., 3rd; Jurecka, P. Refinement of the Cornell et al. Nucleic Acids Force Field Based on Reference Quantum Chemical Calculations of Glycosidic Torsion Profiles. J Chem Theory Comput 2011, 7 (9), 2886–2902. DOI: 10.1021/ct200162x From NLM PubMed-not-MEDLINE.

(43) Steinbrecher, T.; Latzer, J.; Case, D. A. Revised AMBER parameters for bioorganic phosphates. J Chem Theory Comput 2012, 8 (11), 4405–4412. DOI: 10.1021/ct300613v From NLM PubMed-not-MEDLINE.

(44) Izadi, S.; Anandakrishnan, R.; Onufriev, A. V. Building Water Models: A Different Approach. J Phys Chem Lett 2014, 5 (21), 3863–3871. DOI: 10.1021/jz501780a From NLM Publisher.

(45) Mukhopadhyay, A.; Fenley, A. T.; Tolokh, I. S.; Onufriev, A. V. Charge hydration asymmetry: the basic principle and how to use it to test and improve water models. J Phys Chem B 2012, 116 (32), 9776–9783. DOI: 10.1021/jp305226j From NLM Medline.

(46) Cheatham, T. I.; Miller, J.; Fox, T.; Darden, T.; Kollman, P. Molecular dynamics simulations on solvated biomolecular systems: the particle mesh Ewald method leads to stable trajectories of DNA, RNA, and proteins. Journal of the American Chemical Society 1995, 117 (14), 4193–4194.

(47) Berendsen, H. J. C.; Postma, J. P. M.; van Gunsteren, W. F.; DiNola, A.; Haak, J. R. Molecular dynamics with coupling to an external bath. The Journal of Chemical Physics 1984, 81 (8), 3684–3690. DOI: 10.1063/1.448118 (acccessed 7/11/2024).

(48) Kufareva, I.; Ilatovskiy, A. V.; Abagyan, R. Pocketome: an encyclopedia of small-molecule binding sites in 4D. Nucleic Acids Res 2012, 40 (Database issue), D535-540. DOI: 10.1093/nar/gkr825 From NLM Medline.

(49) Yazdani, K.; Jordan, D.; Yang, M.; Fullenkamp, C. R.; Calabrese, D. R.; Boer, R.; Hilimire, T.; Allen, T. E. H.; Khan, R. T.; Schneekloth Jr., J. S. Machine Learning Informs RNA-Binding Chemical Space. Angewandte Chemie International Edition 2023, 62 (11), e202211358. DOI: 10.1002/anie.202211358.

(50) Turbant, F.; Mosca, K.; Busi, F.; Arluison, V.; Wien, F. Circular and Linear Dichroism for the Analysis of Small Noncoding RNA Properties. In Bacterial Regulatory RNA: Methods and Protocols, Arluison, V., Valverde, C. Eds.; Springer US, 2024; pp 399–416.

(51) Siegfried, N. A.; Busan, S.; Rice, G. M.; Nelson, J. A.; Weeks, K. M. RNA motif discovery by SHAPE and mutational profiling (SHAPE-MaP). Nat Methods 2014, 11 (9), 959–965. DOI: 10.1038/nmeth.3029 From NLM Medline.

(52) Nguyen, T. H. D.; Galej, W. P.; Bai, X. C.; Oubridge, C.; Newman, A. J.; Scheres, S. H. W.; Nagai, K. Cryo-EM structure of the yeast U4/U6.U5 tri-snRNP at 3.7 A resolution. Nature 2016, 530 (7590), 298–302. DOI: 10.1038/nature16940 From NLM Medline.

(53) Hardin, J. W.; Warnasooriya, C.; Kondo, Y.; Nagai, K.; Rueda, D. Assembly and dynamics of the U4/U6 di-snRNP by single-molecule FRET. Nucleic Acids Res 2015, 43 (22), 10963–10974. DOI: 10.1093/nar/gkv1011 From NLM Medline.

(54) Aukema, K. G.; Chohan, K. K.; Plourde, G. L.; Reimer, K. B.; Rader, S. D. Small molecule inhibitors of yeast pre-mRNA splicing. ACS Chem Biol 2009, 4 (9), 759–768. DOI: 10.1021/cb900090z From NLM Medline.

(55) Bertram, K.; Agafonov, D. E.; Dybkov, O.; Haselbach, D.; Leelaram, M. N.; Will, C. L.; Urlaub, H.; Kastner, B.; Luhrmann, R.; Stark, H. Cryo-EM Structure of a Pre-catalytic Human Spliceosome Primed for Activation. Cell 2017, 170 (4), 701–713 e711. DOI: 10.1016/j.cell.2017.07.011 From NLM Medline.

(56) Bali, S. K.; Marion, A.; Ugur, I.; Dikmenli, A. K.; Catak, S.; Aviyente, V. Activity of Topotecan toward the DNA/Topoisomerase I Complex: A Theoretical Rationalization. Biochemistry 2018, 57 (9), 1542–1551. DOI: 10.1021/acs.biochem.7b01297 From NLM Medline.

(57) Bocian, W.; Kawecki, R.; Bednarek, E.; Sitkowski, J.; Williamson, M. P.; Hansen, P. E.; Kozerski, L. Binding of topotecan to a nicked DNA oligomer in solution. Chemistry 2008, 14 (9), 2788–2794. DOI: 10.1002/chem.200700732 From NLM Medline.

(58) Rizvi, N. F.; Nickbarg, E. B. RNA-ALIS: Methodology for screening soluble RNAs as small molecule targets using ALIS affinity-selection mass spectrometry. Methods 2019, 167, 28–38. DOI: 10.1016/j.ymeth.2019.04.024 From NLM Medline.

(59) Shortridge, M. D.; Varani, G. Efficient NMR Screening Approach to Discover Small Molecule Fragments Binding Structured RNA. ACS Med Chem Lett 2021, 12 (8), 1253–1260. DOI: 10.1021/acsmedchemlett.1c00109 From NLM PubMed-not-MEDLINE.

(60) Hymon, D.; Martins, J.; Richter, C.; Sreeramulu, S.; Wacker, A.; Ferner, J.; Patwardhan, N. N.; Hargrove, A. E.; Schwalbe, H. NMR (1)H,(19)F-based screening of the four stem-looped structure 5_SL1-SL4 located in the 5’-untranslated region of SARS-CoV 2 RNA. RSC Med Chem 2024, 15 (1), 165–177. DOI: 10.1039/d3md00322a From NLM PubMed-not-MEDLINE.

(61) Haniff, H. S.; Knerr, L.; Chen, J. L.; Disney, M. D.; Lightfoot, H. L. Target-Directed Approaches for Screening Small Molecules against RNA Targets. SLAS Discov 2020, 25 (8), 869–894. DOI: 10.1177/2472555220922802 From NLM Medline.

(62) Cai, Z.; Zafferani, M.; Akande, O. M.; Hargrove, A. E. Quantitative Structure-Activity Relationship (QSAR) Study Predicts Small-Molecule Binding to RNA Structure. J Med Chem 2022, 65 (10), 7262–7277. DOI: 10.1021/acs.jmedchem.2c00254 From NLM Medline.

(63) Velagapudi, S. P.; Seedhouse, S. J.; French, J.; Disney, M. D. Defining the RNA internal loops preferred by benzimidazole derivatives via 2D combinatorial screening and computational analysis. J Am Chem Soc 2011, 133 (26), 10111–10118. DOI: 10.1021/ja200212b From NLM Medline.

(64) Suresh, B. M.; Li, W.; Zhang, P.; Wang, K. W.; Yildirim, I.; Parker, C. G.; Disney, M. D. A general fragment-based approach to identify and optimize bioactive ligands targeting RNA. Proceedings of the National Academy of Sciences 2020, 117 (52), 33197–33203. DOI: doi:10.1073/pnas.2012217117.

(65) Aminova, O.; Disney, M. D. A Microarray-Based Method to Perform Nucleic Acid Selections. In Small Molecule Microarrays: Methods and Protocols, Uttamchandani, M., Yao, S. Q. Eds.; Humana Press, 2010; pp 209–224.

(66) Ursu, A.; Childs-Disney, J. L.; Angelbello, A. J.; Costales, M. G.; Meyer, S. M.; Disney, M. D. Gini Coefficients as a Single Value Metric to Define Chemical Probe Selectivity. ACS Chem Biol 2020, 15 (8), 2031–2040. DOI: 10.1021/acschembio.0c00486 From NLM Medline.

