## Supplementary Information for "Discovery of small molecule probes targeting the 5′ stem-loop in the yeast U4/U6 snRNA"

**in yeast U4/U6 snRNA assembly**

Mo Yang^1^, Shaifaly Parmar^1^, Sumirtha Balaratnam^1^, Desta D. Bume^1^, Peri R. Prestwood^1^, Wojciech K. Kasprzak^2^, and John S. Schneekloth, Jr.^1,^ *

^1^Chemical Biology Laboratory, Center for Cancer Research, National Cancer Institute, Frederick, MD 21702-1201, USA

^2^Advanced Biomedical Computational Science, Frederick National Laboratory for Cancer Research, Frederick, MD 21702, USA

**Supplementary Tables**

**Table S1.** Pocket analysis of U4/U6 snRNA assembly

| **No.** | **Volume** | **Hydrophobicity** | **Buriedness** | **Aromatic** | **DLID** | **Area** | **Radius** | **Nonsphericity** |
| --- | --- | --- | --- | --- | --- | --- | --- | --- |
| 1 | 1131 | 0.334 | 0.794 | 0.026 | 0.378 | 1011 | 6.464 | 1.926 |
| **2** | **449.9** | **0.539** | **0.774** | **0.172** | **0.0835** | **389.2** | **4.753** | **1.371** |
| **3** | **352.8** | **0.446** | **0.738** | **0.188** | **-0.45** | **335** | **4.383** | **1.387** |
| 4 | 188.5 | 0.352 | 0.739 | 0.0472 | -1.12 | 256.4 | 3.557 | 1.613 |
| 5 | 184.5 | 0.449 | 0.711 | 0.0176 | -1.03 | 205.3 | 3.531 | 1.31 |
| 6 | 122.7 | 0.325 | 0.657 | 0 | -1.82 | 201.2 | 3.082 | 1.685 |
| 7 | 105.4 | 0.476 | 0.633 | 0.115 | -1.69 | 155.4 | 2.93 | 1.44 |
| 8 | 112.7 | 0.467 | 0.703 | 0 | -1.38 | 156.3 | 2.997 | 1.385 |
| 9 | 100.1 | 0.602 | 0.740 | 0.0468 | -1.02 | 134.4 | 2.88 | 1.29 |
| 10 | 108.5 | 0.565 | 0.754 | 0.00431 | -0.986 | 134.8 | 2.958 | 1.225 |

**Table S2.** U4/U6 snRNA constructs used in SMM screening.

| **No.** | **Sequence (5′-3')** | **Length** |
| --- | --- | --- |
| 1 | GGC CUU AUG CAC GGG AAA UAC GCA UAU CAG UGA GGA UUC GUC CGA GAU UGU GUU UUU GCU GGU GUA AAU CAG CAG UUC CCC UGC AUA AGG CU | 92 nt |
| 2 | GGC CUU AUG CAC GGG AAA UAC GCA UAU CGU AAG AUU GUG UUU UUG CUG GUG UAA AUC AGC AGU UCC CCU GCA UAA GGC U | 79 nt |
| 3 | GGG AAA UAC GCA UAU CAG UGA GGA UUC GUC CGA GAU UGU GUU UUU GCU GGU GUA AAU CAG CAG UUC CC | 68 nt |
| 4 | CGC AUA UCA GUG AGG AUU CGU CCG AGA UUG UG | 32 nt |

**Table S3.** Sequence of oligos and primers used in the study.

|  | **Sequence (5′-3')** |
| --- | --- |
| Template | TAATACGACTCACTATAGGGCCTTCGGGCCAAGGCCTTATGCACGGGAAATACGCATATCAGTGAGGATTCGTCCGAGATTGTGTTTTTGCTGGTGTAAATCAGCAGTTCCCCTGCATAAGGCTTCGATCCGGTTCGCCGGATCCAAATCGGGCTTCGGTCCGGTTC |
| Template_F | TAATACGACTCACTATAGGGCC |
| Template_R | GAACCGGACCGAAGCC |
| RT oligo | GAACCGGACCGAAGCCCG |
| Step1_F: | GACTGGAGTTCAGACGTGTGCTCTTCCGATCTNNNNNGGCCTTATGCACGGGA |
| Step1_R | CCCTACACGACGCTCTTCCGATCTNNNNNGAACCGGACCGAAGCCCG |
| Step2_F | CAAGCAGAAGACGGCATACGAGATTCGCCTTAGTGACTGGAGTTCAGAC |
| Step2_R | AATGATACGGCGACCACCGAGATCTACACTCTTTCCCTACACGACGCTCTTCCG |

**Supplementary Figures**

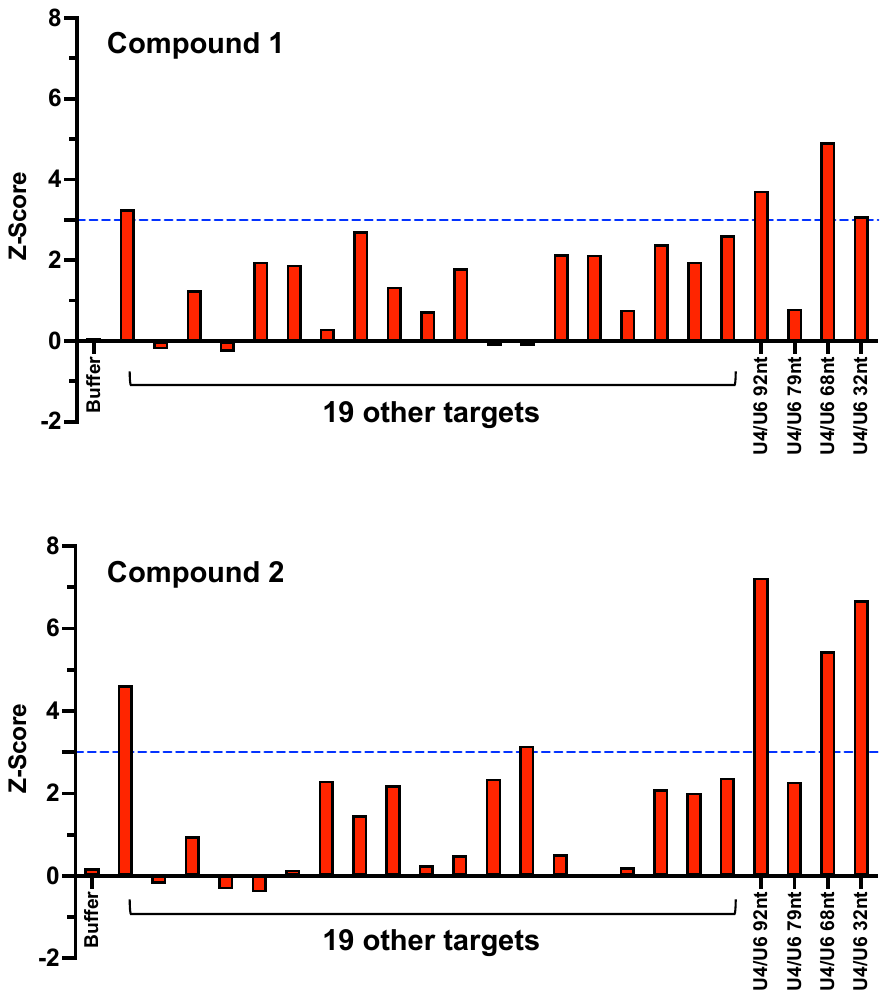

**Figure S1.** Selectivity profiling of two hit compounds (**1** and **2**) based on SMM Z-scores in the compiled dataset. 19 other targets (DNA/RNA structures, list not shown) that were previously screened are used for comparison.

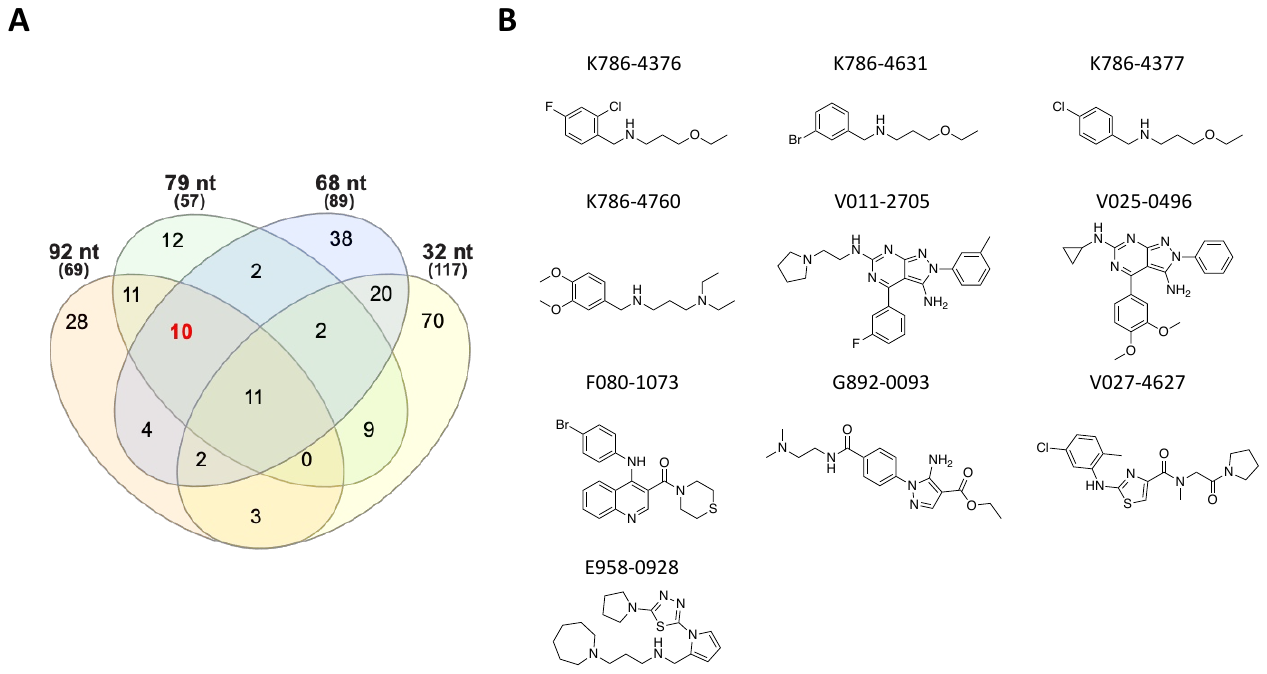

**Figure S2.** (A) Ten hit compounds that potentially bind 3WJ (positive in screens using 92 nt & 79 nt & 68 nt RNAs). (B) Chemical structures of the compounds and their commercial IDs in ChemDiv library.

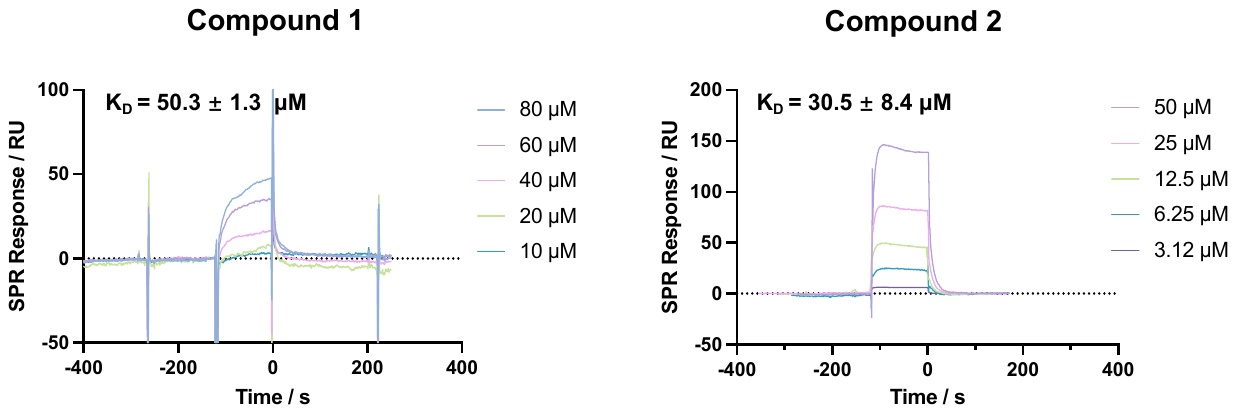

**Figure S3.** Binding affinity measurement of two hits (compounds **1** and **2**) by SPR. U4/U6 92-nt RNA constructed was 5′-biotin-labeled and immobilized on the SPR chip surface.

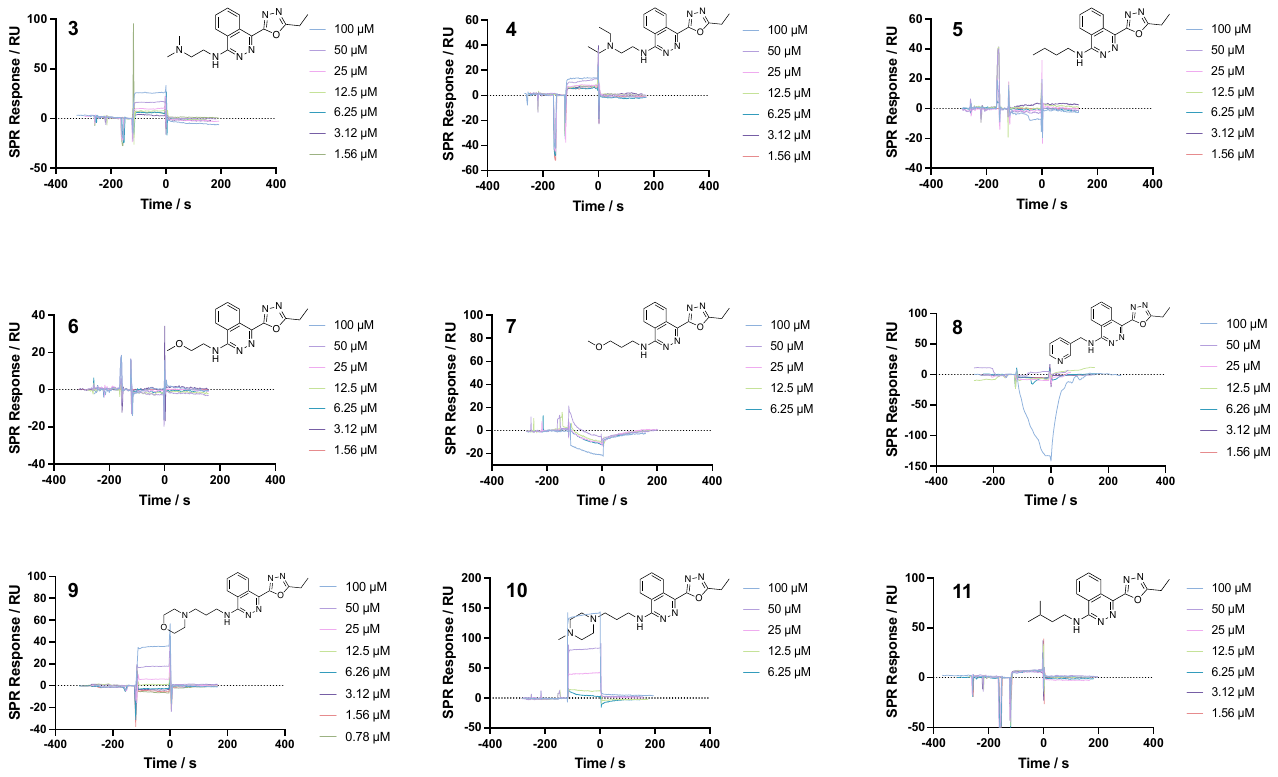

**Figure S4-1.** SPR curves of hit **2** analogs and binding affinity measurement. SPR curves show titrations using **3**-**11**.

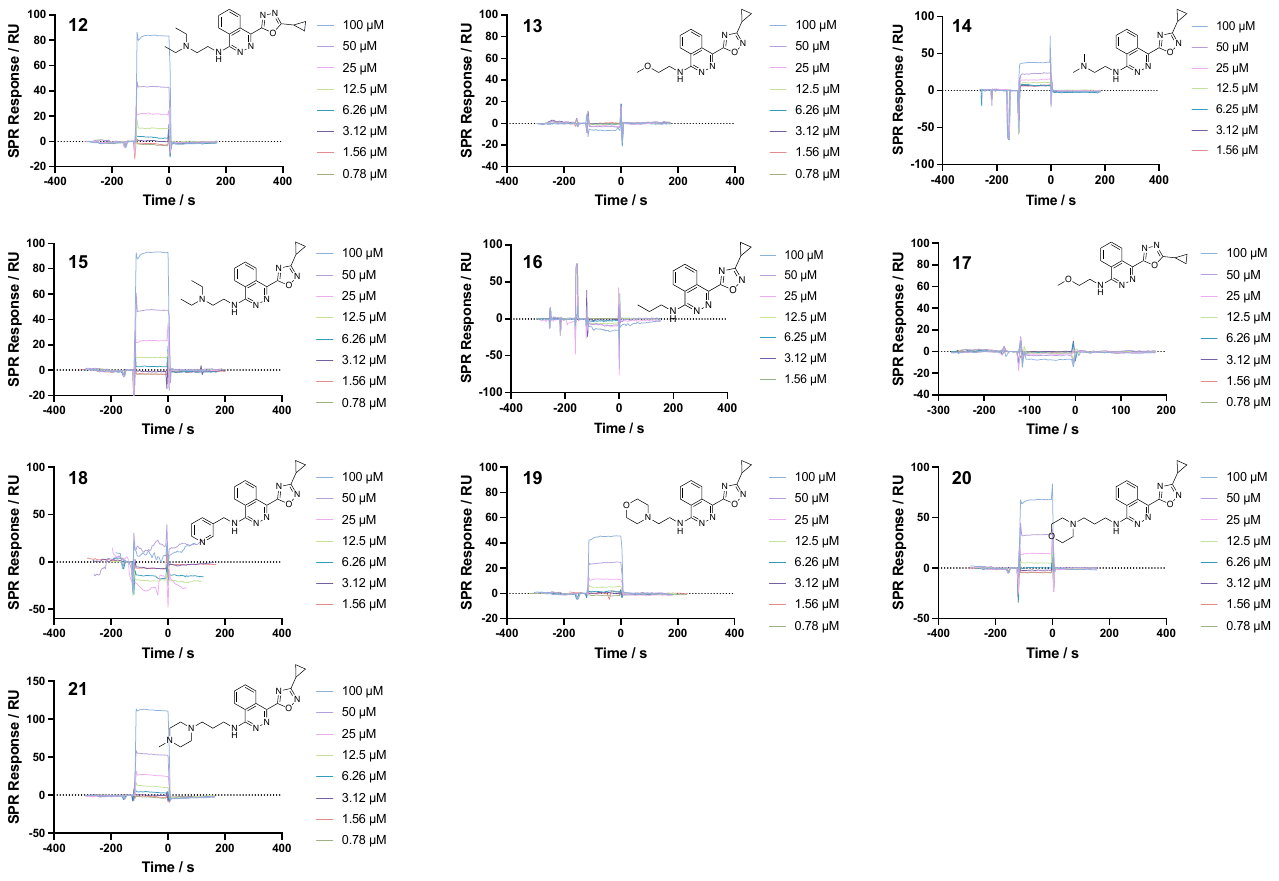

**Figure S4-2.** SPR curves of hit **2** analogs and binding affinity measurement. SPR curves show titrations using **12**-**21**.

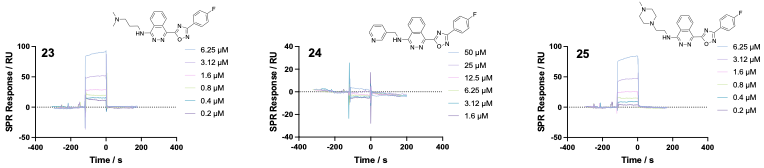

**Figure S4-3.** SPR curves of hit **2** analogs and binding affinity measurement. SPR curves show titrations using **23**-**25**.

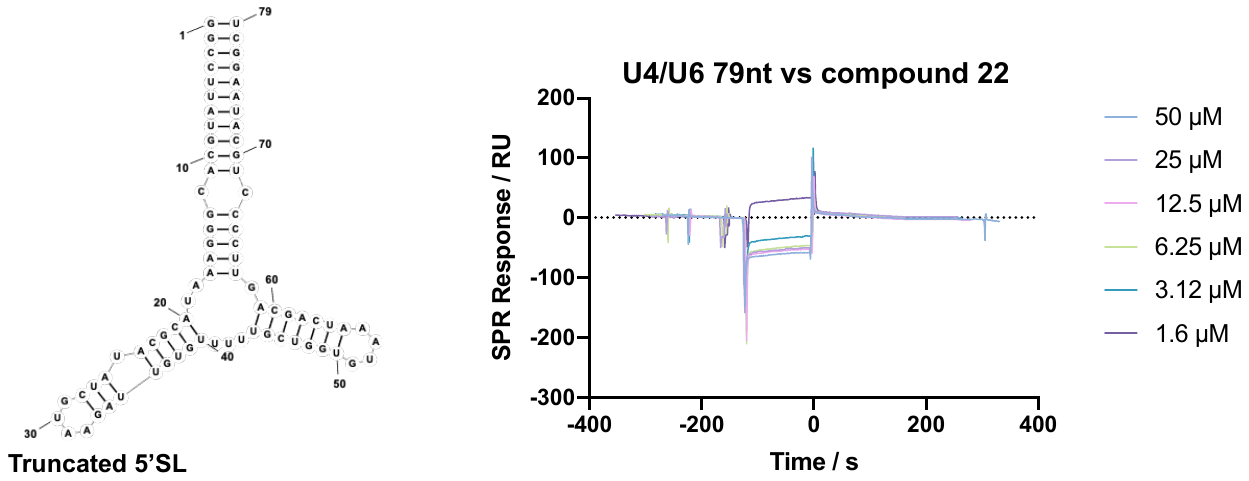

**Figure S5.** The SPR binding test between the optimized compound 22 and U4/U6 79-nt snRNA construct lacks a K-turn motif.

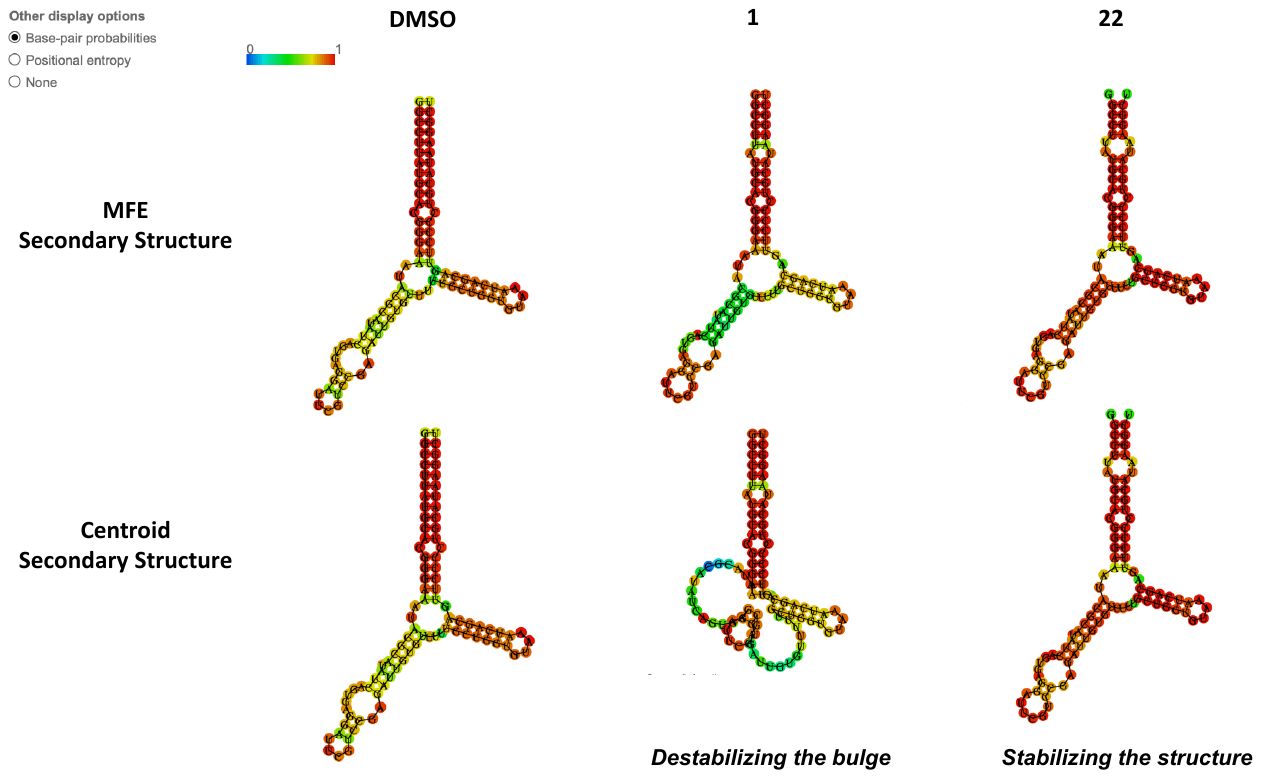

**Figure S6.** SPR titration curves of all the analogs and binding affinity measurement. The results of RNA secondary structure prediction (MFE and Centroid) are visualized by RNAfold Webserver (http://rna.tbi.univie.ac.at/cgi-bin/RNAWebSuite/RNAfold.cgi).

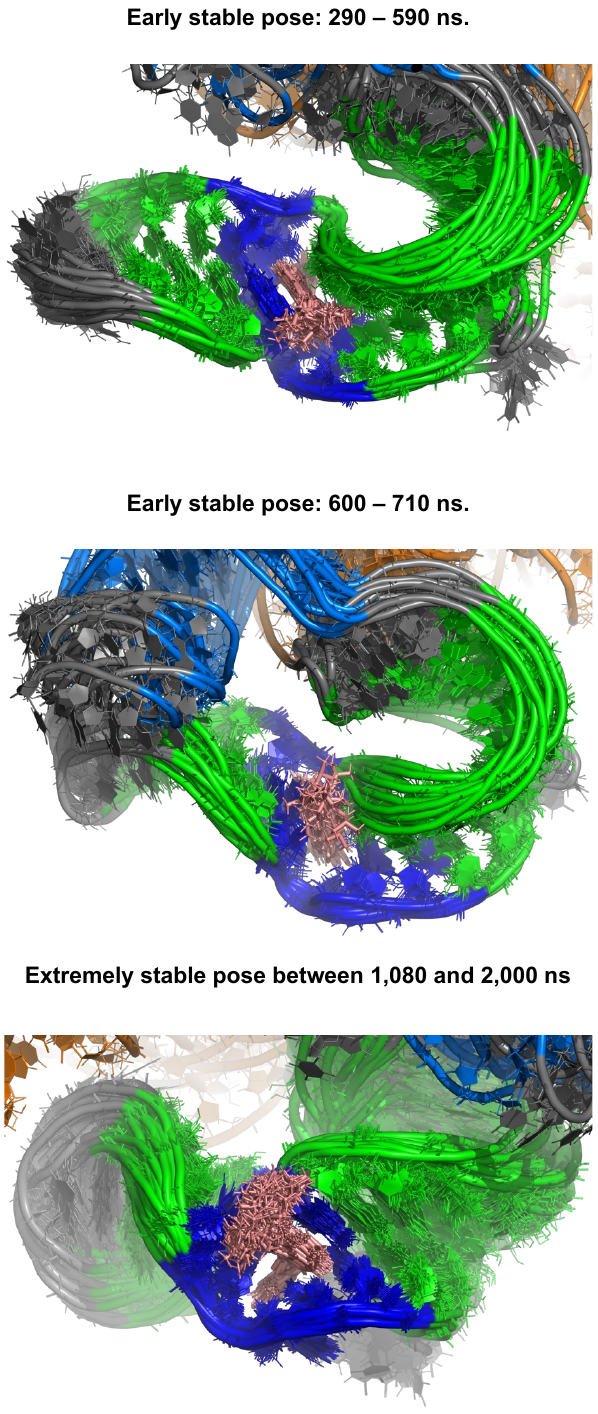

**Figure S7.** Poses of simulated RNA-compound **22** complex demonstrating intermediate states during MD simulation.

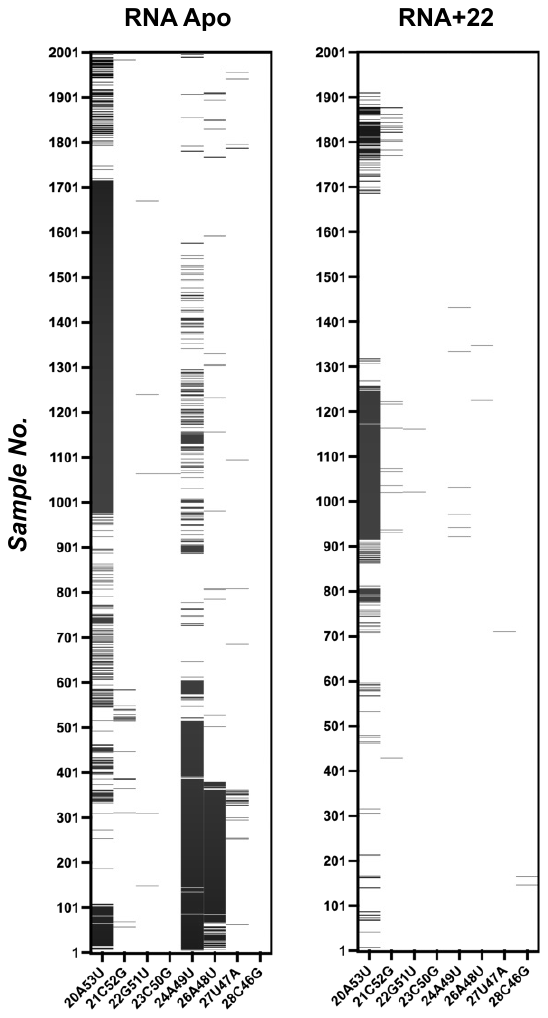

**Figure S8.** A sampling of base pairing status in duplex regions during 2000 ns simulation. A total of 2000 samples were calculated; white = paired; and black = unpaired.

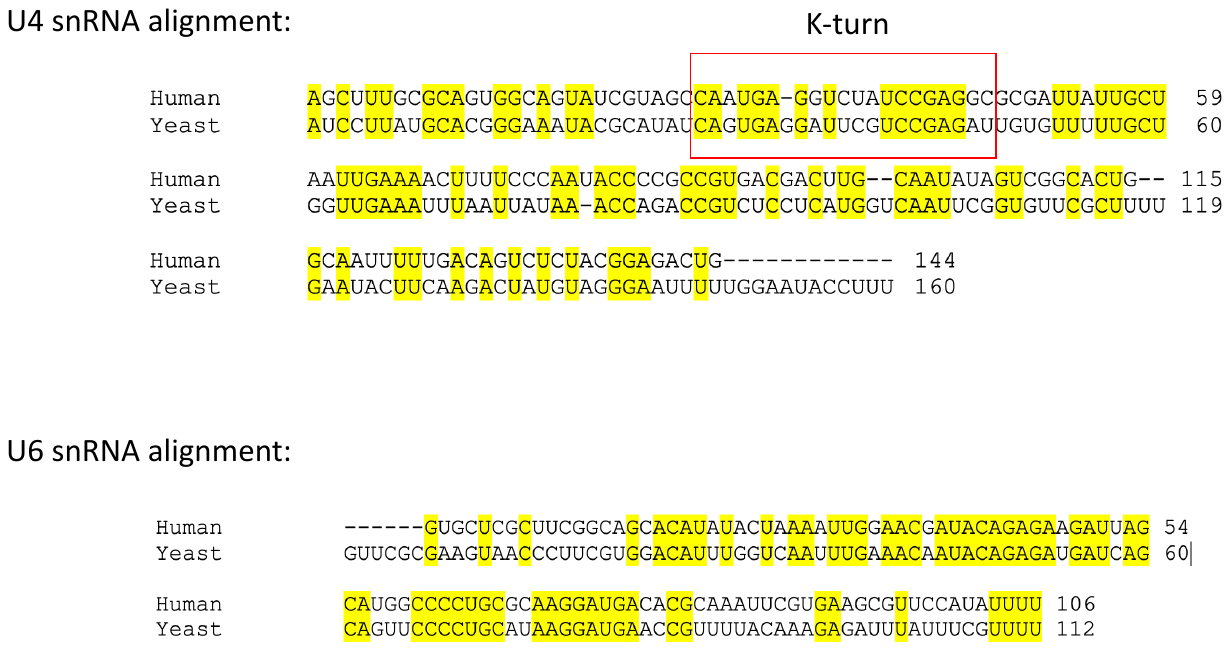

**Figure S9.** Sequence alignment and comparison of yeast and human U4/U6 snRNAs.

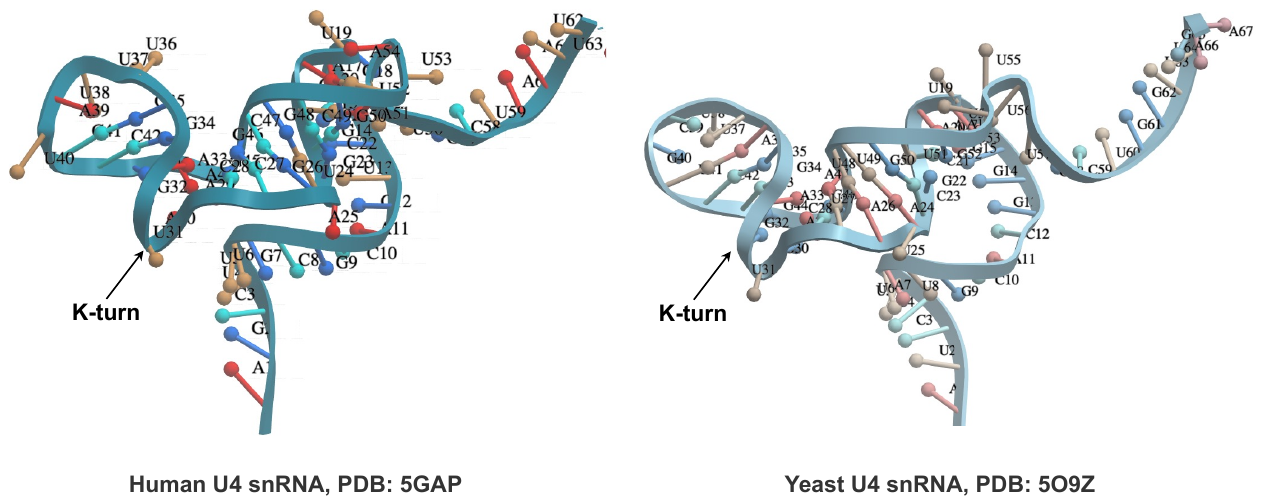

**Figure S10.** Side-by-side comparison of 5′ SL between human and yeast U4 snRNA. The structural information was obtained from published data solved by Cryo-EM.^1, 2^ Both structures are shown at protein-binding states in the snRNP complex while the proteins are hidden.

**Chemical synthesis and NMR data**

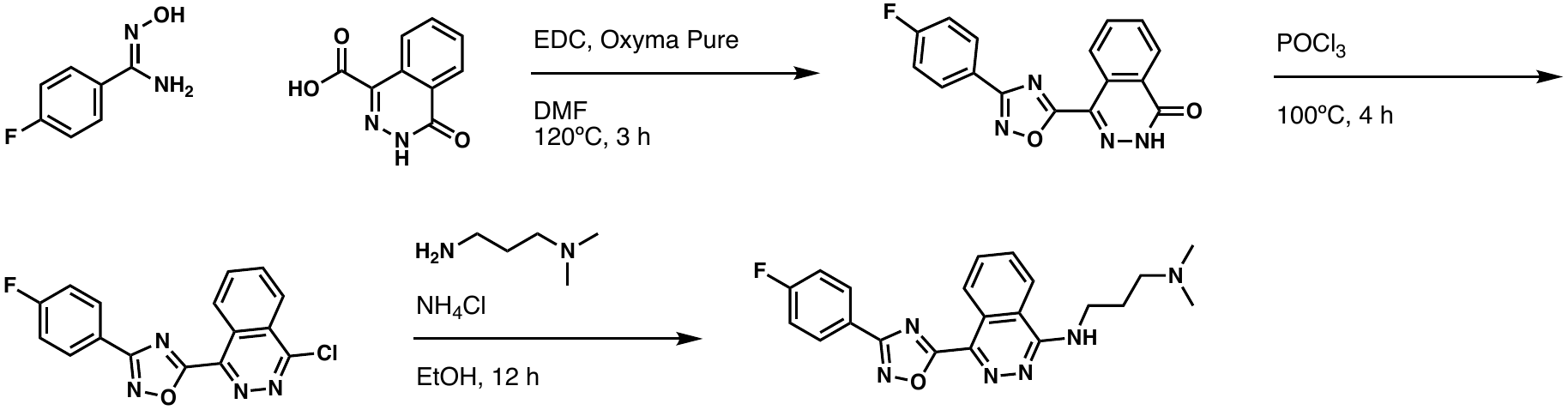

**23** (273C-027)

A solution of 4-oxo-3,4-dihydrophthalazine-1-carboxylic acid (500 mg, 2.6 mmol), EDC (450 mg, 2.9 mmol), and Oxyma Pure (444 mg, 2.9 mmol) in DMF (10 mL) was stirred at room temperature for 10 minutes, then (*Z*)-4-fluoro-*N*'-hydroxybenzimidamide (526 mg, 3.4 mmol) was added. The mixture was heated to 120ºC and stirred for three hours. After cooling to room temperature, the reaction was quenched and diluted with water. The crude product was filtered and dried, then taken to the next step.

To 4-(3-(4-fluorophenyl)-1,2,4-oxadiazol-5-yl)phthalazin-1(2*H*)-one (150 mg, 0.49 mmol) in a round bottom flask was added POCl_3_ (3 mL). The mixture was heated to 100ºC and stirred for two hours. The reaction was then cooled to room temperature, poured into ice-cold water, and neutralized with 10% NaOH. The organic layer was extracted using EtOAc (3 x 10 mL), washed with brine (10 mL), then dried (Na_2_SO_4_). The crude product was purified by ISCO flash column chromatography (MeOH:DCM, 0-30%).

A solution of 5-(4-chlorophthalazin-1-yl)-3-(4-fluorophenyl)-1,2,4-oxadiazole (20 mg, 0.060 mmol) and *N*^1^,*N*^1^-dimethylpropane-1,3-diamine (30 µL, 0.24 mmol) in EtOH (2 mL) was heated to 80ºC. The reaction stirred for 12 hours, then was returned to room temperature and concentrated down in vacuo. The crude product was purified by ISCO flash column chromatography (MeOH:DCM, 0-30%) to afford the desired product.

^1^H NMR (400 MHz, DMSO-*d*_6_): δ 9.11 (dd, *J* = 8.2, 0.9 Hz, 1H), 8.80 (t, *J* = 5.7 Hz, 1H), 8.58 (d, *J* = 8.2 Hz, 1H), 8.23-8.18 (m, 2H), 8.04-7.99 (m, 1H), 7.98-7.93 (m, 1H), 7.47-7.41 (m, 2H), 3.78 (q, *J* = 6.3 Hz, 2H), 3.10 (t, *J* = 7.8 Hz, 2H), 2.68 (s, 6H), 2.16-2.08 (m, 2H); ^13^C NMR (125 MHz, DMSO-*d*_6_): δ 173.3, 167.1, 165.3-162.8 (d, *J^1^_C-F_* = 248.0 Hz), 154.8, 136.1, 133.1, 131.8, 129.8-129.7 (d, *J^3^_C-F_* = 8.8 Hz), 124.8, 124.7, 123.0, 122.8 (d, *J^4^_C-F_* = 2.6 Hz), 116.8, 116.6-116.4 (d, *J^2^_C-F_* = 22.3 Hz), 54.7, 42.2, 40.4, 38.4, 23.6; LCMS: (ESI+) m/z calculated for C_21_H_21_FN_6_O [M+H]^+^: 393.18, found: 393.40, rt: 3.78 min.

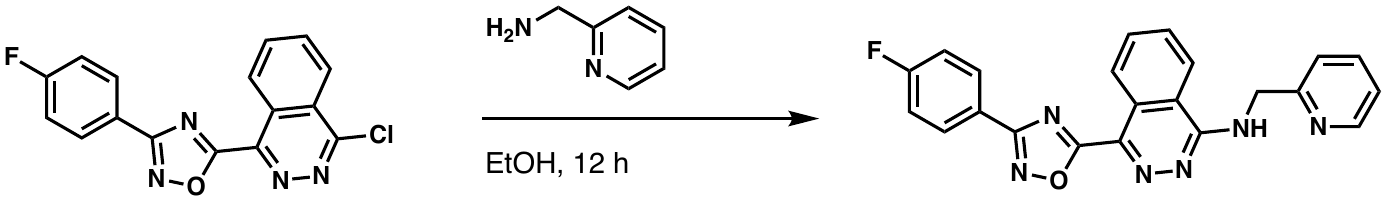

**24** (273B-203)

A solution of 5-(4-chlorophthalazin-1-yl)-3-(4-fluorophenyl)-1,2,4-oxadiazole (30 mg, 0.090 mmol) and pyridin-2-ylmethanamine (18 µL, 0.18 mmol) in EtOH (3 mL) was heated to 80ºC. The reaction stirred for 12 hours, then was returned to room temperature and concentrated down in vacuo. The crude product was purified by ISCO flash column chromatography (MeOH:DCM, 0-30%) to afford the desired product.

^1^H NMR (400 MHz, DMSO-*d*_6_): δ 9.20 (dd, *J* = 8.5, 1.2 Hz, 1H), 8.99 (t, *J* = 5.8 Hz, 1H), 8.70 (d, *J* = 1.3 Hz, 1H), 8.50 (d, *J* = 8.0 Hz, 1H), 8.47 (dd, *J* = 4.7, 1.4 Hz, 1H), 8.28-8.23 (m, 2H), 8.12-8.03 (m, 2H), 7.87-7.83 (m, 1H), 7.52-7.46 (m, 2H), 7.37 (dd, *J* = 7.9, 4.6 Hz, 1H), 4.98 (d, *J* = 5.9 Hz, 2H);

^13^C NMR (125 MHz, DMSO-*d*_6_): δ 175.6, 167.2, 165.2-162.5 (d, *J^1^_C-F_* = 268.8 Hz), 154.4, 149.1, 148.1, 139.4, 136.7, 135.3, 133.2, 133.1, 129.9-129.8 (d, *J^3^_C-F_* = 9.0 Hz), 128.4, 125.0, 123.5, 122.6, 116.8, 116.7-116.4 (d, *J^2^_C-F_* = 22.4 Hz), 110.3, 42.2; LCMS: (ESI+) m/z calculated for C_22_H_15_FN_6_O [M+H]^+^: 399.14, found: 399.35, rt: 3.88 min.

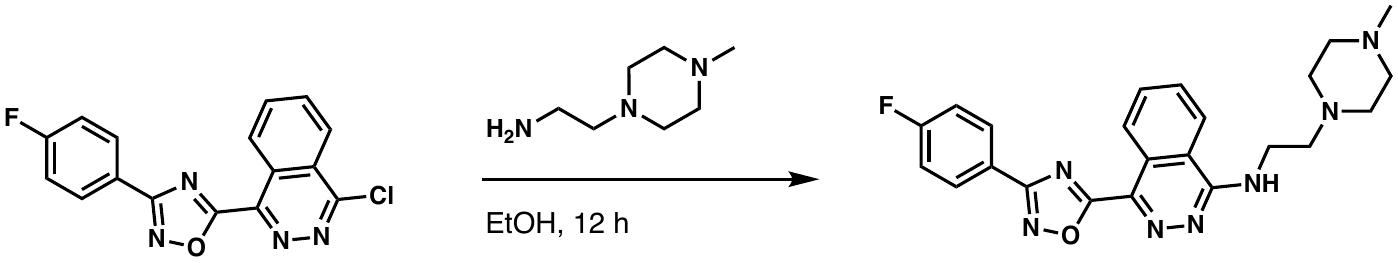

**25** (273C-028)

A solution of 5-(4-chlorophthalazin-1-yl)-3-(4-fluorophenyl)-1,2,4-oxadiazole (20 mg, 0.060 mmol) and 2-(4-methylpiperazin-1-yl)ethan-1-amine (38 µL, 0.24 mmol) in EtOH (3 mL) was heated to 80ºC. The reaction stirred for 12 hours, then was returned to room temperature and concentrated down in vacuo. The crude product was purified by ISCO flash column chromatography (MeOH:DCM, 0-30%) to afford the desired product.

^1^H NMR (400 MHz, DMSO-*d*_6_): δ 9.09 (dd, *J* = 8.5, 1.2 Hz, 1H), 8.37 (d, *J* = 7.8 Hz, 1H), 8.29 (t, *J* = 5.6 Hz, 1H), 8.20-8.15 (m, 2H), 8.02-7.97 (m, 1H), 7.96-7.91 (m, 1H), 7.45-7.39 (m, 2H), 3.79 (q, *J* = 6.5 Hz, 2H), 2.64 (t, *J* = 6.9 Hz, 2H), 2.47-2.44 (m, 4H), 2.36-2.26 (m, 4H), 2.12 (s, 3H); ^13^C NMR (125 MHz, DMSO-*d*_6_): δ 173.3, 167.1, 165.3-162.8 (d, *J^1^_C-F_* = 248.0 Hz), 154.6, 135.9, 133.0, 131.8, 129.8-129.7 (d, *J^3^_C-F_* = 8.7 Hz), 124.8, 122.8 (d, *J^4^_C-F_* = 2.6 Hz), 122.5, 116.6, 116.6-116.4 (d, *J^2^_C-F_* = 22.1 Hz), 56.2, 54.7, 52.7, 45.6, 40.4; LCMS: (ESI+) m/z calculated for C_23_H_24_FN_7_O [M+H]^+^: 434.21, found: 434.60, rt: 3.69 min.

**NMR Spectra**

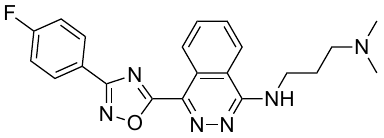

**23**

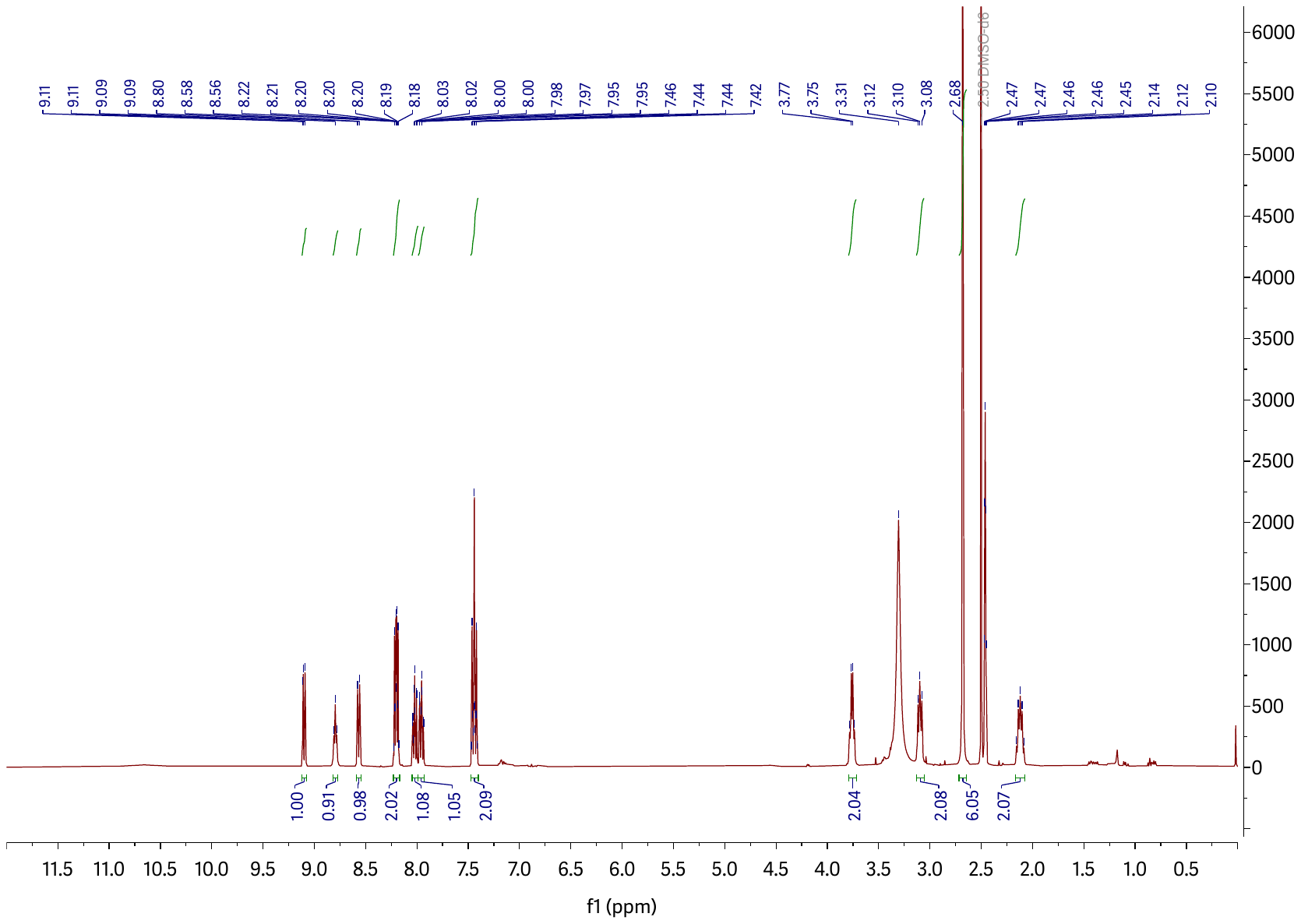

**23 ^1^H NMR.** 400 MHz, DMSO-*d*_6_

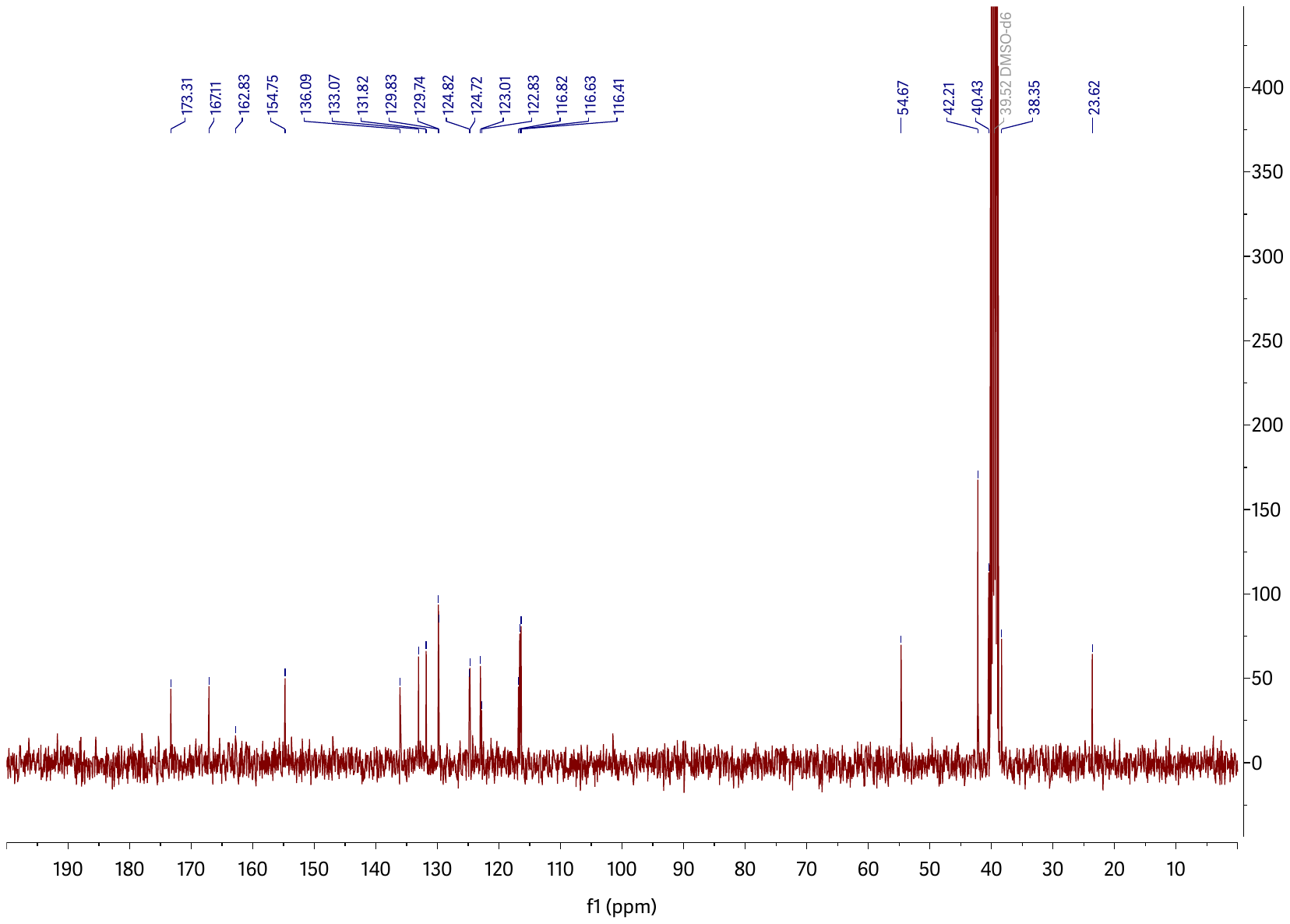

**23 ^13^C NMR.** 125 MHz, DMSO-*d*_6_

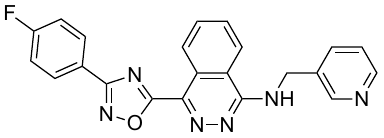

**24**

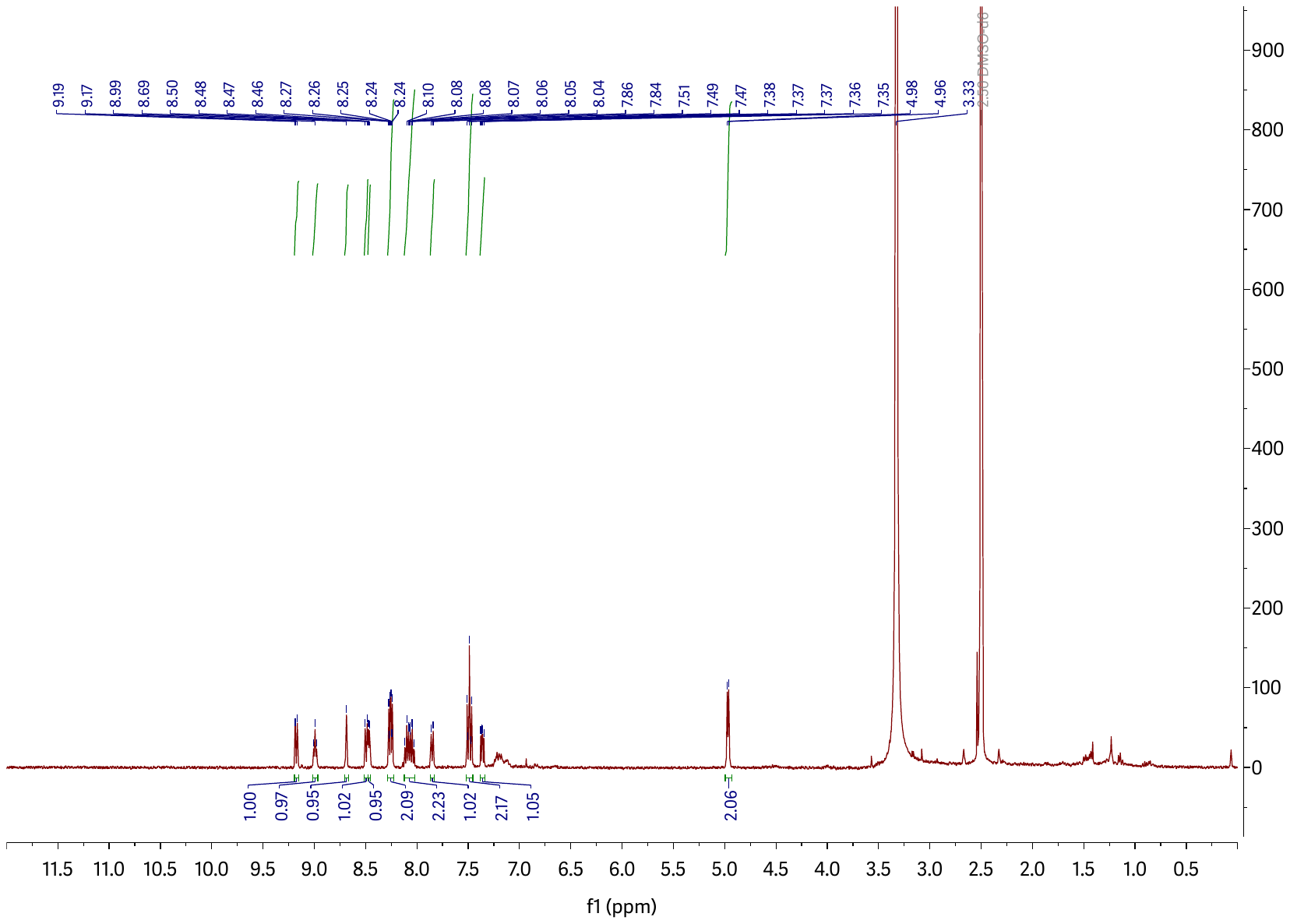

**24 ^1^H NMR.** 400 MHz, DMSO-*d*_6_

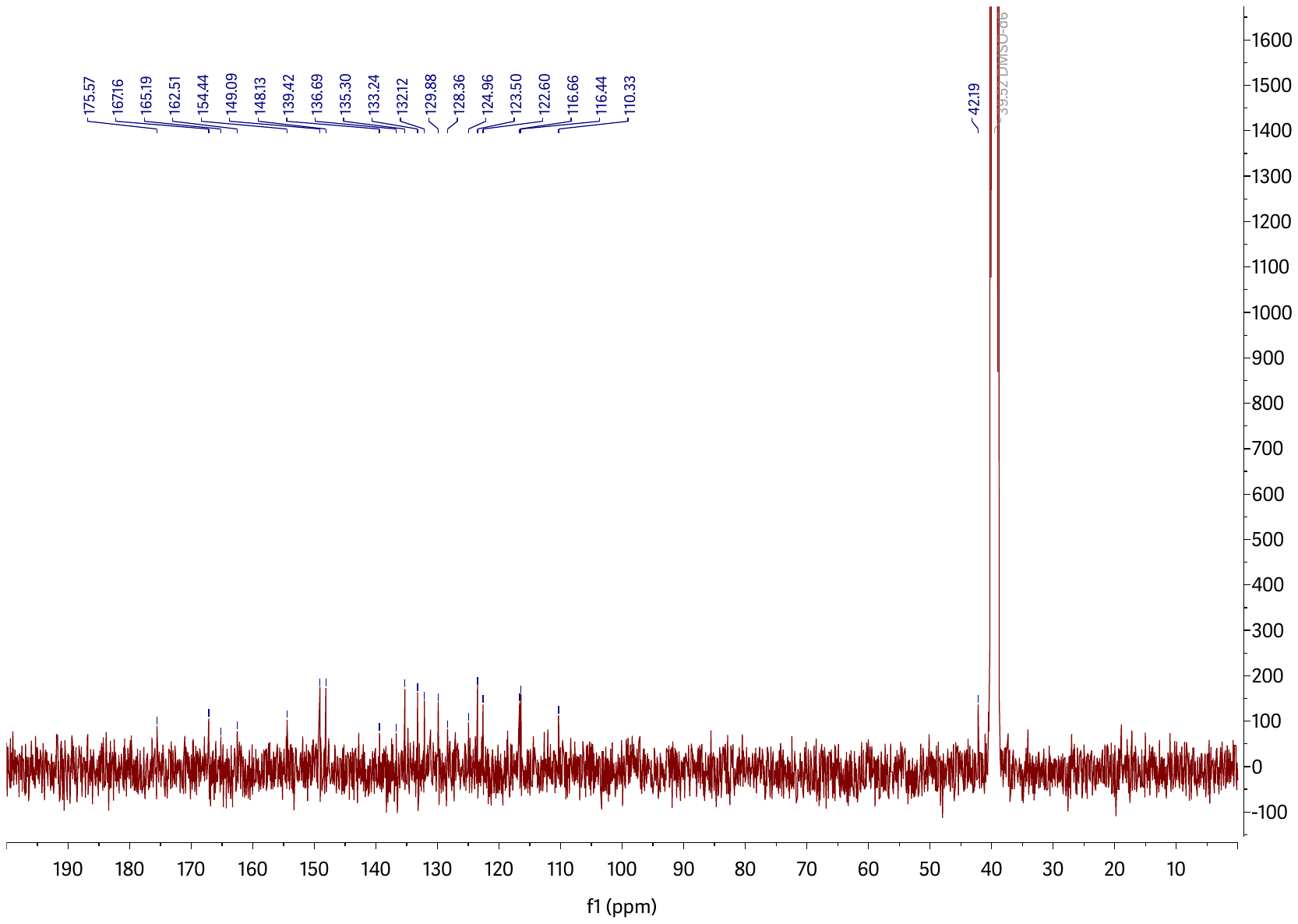

**24 ^13^C NMR.** 125 MHz, DMSO-*d*_6_

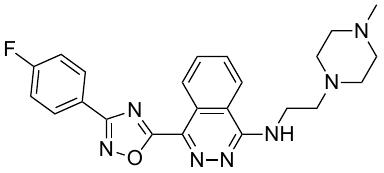

**25**

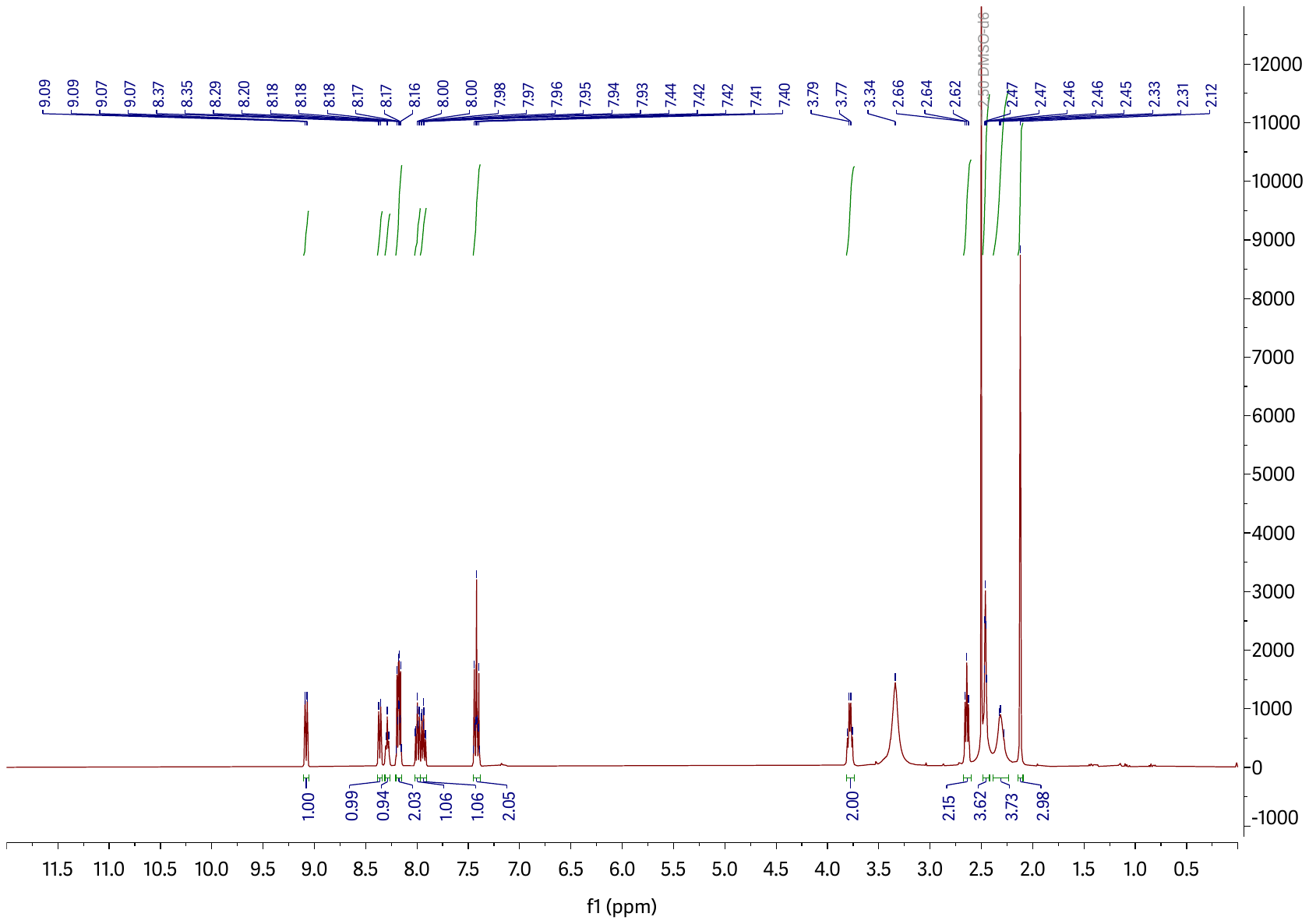

**25 ^1^H NMR.** 400 MHz, DMSO-*d*_6_

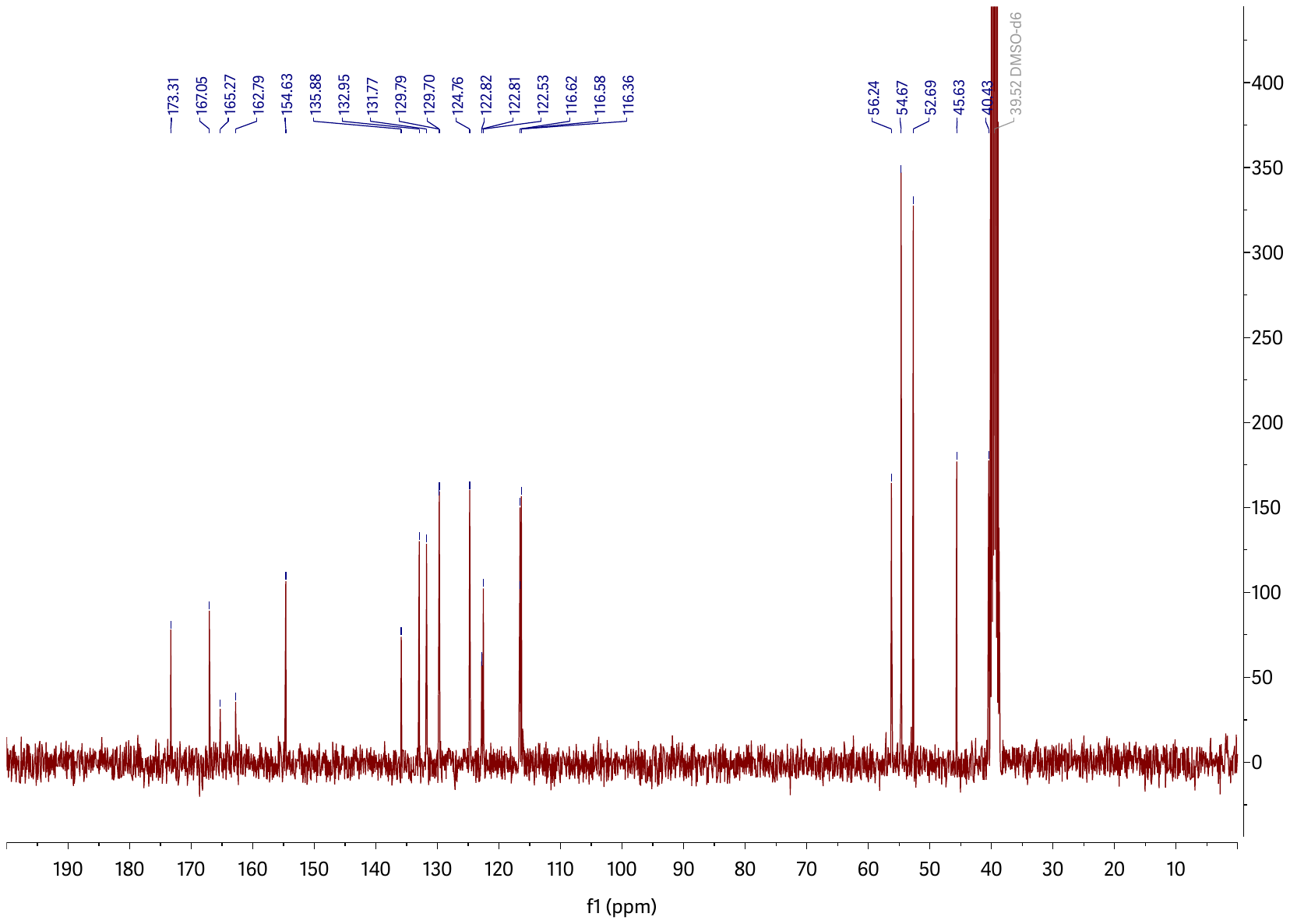

**25 ^13^C NMR.** 125 MHz, DMSO-*d*_6_

**Reference**

(1) Bertram, K.; Agafonov, D. E.; Dybkov, O.; Haselbach, D.; Leelaram, M. N.; Will, C. L.; Urlaub, H.; Kastner, B.; Luhrmann, R.; Stark, H. Cryo-EM Structure of a Pre-catalytic Human Spliceosome Primed for Activation. *Cell* **2017**, *170* (4), 701-713 e711. DOI: 10.1016/j.cell.2017.07.011 From NLM Medline.

(2) Nguyen, T. H. D.; Galej, W. P.; Bai, X. C.; Oubridge, C.; Newman, A. J.; Scheres, S. H. W.; Nagai, K. Cryo-EM structure of the yeast U4/U6.U5 tri-snRNP at 3.7 A resolution. *Nature* **2016**, *530* (7590), 298-302. DOI: 10.1038/nature16940 From NLM Medline.
